# Taxon-specific phytoplankton growth, nutrient utilization, and light limitation in the oligotrophic Gulf of Mexico

**DOI:** 10.1101/2021.03.01.433426

**Authors:** Natalia Yingling, Thomas B. Kelly, Taylor A. Shropshire, Michael R. Landry, Karen E. Selph, Angela N. Knapp, Sven A. Kranz, Michael R. Stukel

## Abstract

The highly stratified, oligotrophic regions of the oceans are predominantly nitrogen limited in the surface ocean and light limited at the deep chlorophyll maximum (DCM). Hence, determining light and nitrogen co-limitation patterns for diverse phytoplankton taxa is crucial to understanding marine primary production throughout the euphotic zone. During two cruises in the deep-water Gulf of Mexico, we measured primary productivity (H^13^CO_3_^−^), nitrate uptake (^15^NO_3_^−^), and ammonium uptake (^15^NH_4_^+^) throughout the water column. Primary productivity declined with depth from the mixed-layer to the DCM, averaging 27.1 mmol C m^−2^ d^−1^. The fraction of growth supported by NO_3_^−^ was consistently low, with upper euphotic zone values ranging from 0.01 to 0.14 and lower euphotic zone values ranging from 0.03 to 0.44. Nitrate uptake showed strong diel patterns (maximum during the day), while ammonium uptake exhibited no diel variability. To parameterize taxon-specific phytoplankton nutrient and light utilization, we used a data assimilation approach (Bayesian Markov Chain Monte Carlo) including primary productivity, nutrient uptake, and taxon-specific growth rate measurements. Parameters derived from this analysis define distinct niches for five phytoplankton taxa (*Prochlorococcus, Synechococcus*, diatoms, dinoflagellates, and prymnesiophytes) and may be useful for constraining biogeochemical models of oligotrophic open-ocean systems.

## INTRODUCTION

Nutrient acquisition by phytoplankton is an important factor regulating marine primary productivity (Davey *et al*., 2008; Duce *et al*., 2008; Mulholland and Lomas, 2008). The open-ocean Gulf of Mexico (GoM) is a highly-stratified, oligotrophic region where the Loop Current, mesoscale eddies and episodic storm events influence lateral and vertical transport of nutrients and organisms (Biggs, 1992; Forristall *et al*., 1992; Biggs and Ressler, 2001; Oey *et al*., 2005). These features create dynamic ecological mosaics with substantial mesoscale spatial variability in the phytoplankton community (Biggs and Müller-Karger, 1994; Gomez *et al*., 2018). Intense stratification also leads to deep chlorophyll maxima (DCM) that potentially provide unique niches for phytoplankton compared to low-nutrient, high-light environments found in the shallow mixed-layer (Shropshire *et al*., 2020; Knapp *et al*., this issue; Selph *et al*., this issue). Additionally, the oligotrophic GoM is an important spawning region for economically-valuable and environmentally-important nekton, including Atlantic Bluefin Tuna (Rooker *et al*., 2007; Cornic *et al*., 2008). In addition to vertical advection, previous studies have found that nitrogen fixation, NO_3_^−^ upwelling along the boundaries of mesoscale eddies, and horizontal advection of nutrients can be important nitrogen sources in the GoM (Walker *et al*., 2005; Mulholland *et al*., 2006). While the sources of bioavailable nitrogen in the oligotrophic GoM are likely spatiotemporally variable, a more accurate understanding of phytoplankton nutrient-uptake dynamics is important to provide insight into what regulates group-specific community composition and inform future biogeochemical models.

Variability in phytoplankton ecophysiology adds complexity to the basic processes of primary productivity in pelagic habitats. *Prochlorococcus* is often the dominant phytoplankton taxon in oligotrophic regions (Chisholm *et al*., 1988; Partensky *et al*., 1999); however, many ecotypes of this genus exist, with different nutrient uptake capabilities and consequently biogeochemical impacts (Zwirglmaier et al., 2008; Martiny *et al*., 2009; Kashtan *et al*., 2014; Kent *et al*., 2016; De Martini *et al*., 2018). *Prochlorococcus* is the numerically dominant phytoplankton in our study region and comprises roughly half of the total carbon-based biomass (Selph *et al*., this issue). *Synechococcus* and picoeukaryotes (especially prymnesiophytes, chlorophytes, and pelagophytes) are also important, with picoeukaryotes comprising a larger portion of biomass with increasing depth (Selph *et al*., this issue). Since different phytoplankton groups, as well as species and ecotypes within groups, differ in their adaptations for life in nutrient-depleted environments, their responses to dynamic environments may also be quantitatively and qualitatively different (Dutkiewicz *et al*., 2013).

Constraining phytoplankton functional relationships (i.e., phytoplankton responses to environmental variables) is necessary to enable forecasting of ecological impacts in dynamic environments and modified physical conditions—such as those expected in a future ocean. Numerical modeling of marine microbial ecosystems is presently limited by an inability to accurately constrain *in situ* ecophysiological relationships amongst diverse phytoplankton taxa (Anderson, 2005; Franks, 2009; Follows and Dutkiewicz, 2011). Here, we address three questions aiming to better constrain group-specific functional phytoplankton relationships: 1) How do net primary production (NPP) and NO_3_^−^ uptake rates of phytoplankton in the open-ocean GoM vary with depth and time of day; 2) What nitrogen sources support these phytoplankton; and 3) How do nutrient limitation and light limitation vary among different phytoplankton taxa?

Answers to these questions are derived from measurements collected during two field studies as part of the Bluefin Larvae in Oligotrophic Ocean Foodwebs: Investigating Nutrients to Zooplankton in the Gulf of Mexico (BLOOFINZ-GoM) project (Gerard *et al*., this issue). Specifically, we conducted Lagrangian experiments with repeated depth-resolved measurements of NPP, nutrient uptake and taxon-specific phytoplankton growth rates, while also assessing the biomass of different phytoplankton. We then assimilate this field data to parameterize group-specific phytoplankton nutrient kinetics for application to biogeochemical models.

## METHODS

### Cruise structure and sampling strategy

Data are from two cruises in the open-ocean northern GoM (NF1704 in May, 2017; NF1802 in May, 2018) as part of the BLOOFINZ-GoM project. We conducted five Lagrangian experiments (‘Cycles’ referred to as C1, C2, C3, C4 and C5 to represent each cycle) lasting 2-4 days each (Gerard *et al*., this issue). Each cycle used a pair of satellite-tracked marker buoys tethered to subsurface drogues to follow the mixed-layer in a Lagrangian frame of reference (Landry *et al*., 2009; Stukel *et al*., 2015). One array (“incubation array”) included attachment points for mesh bags containing incubation bottles for NO_3_^−^ uptake, NPP, and taxon-specific phytoplankton growth and grazing rates at depths spanning the euphotic zone. The Lagrangian approach permitted repeated sampling of the same water parcel to quantify variability in phytoplankton biomass and rates over the duration of each cycle. Samples for fluorometric chlorophyll *a* (Strickland and Parsons, 1972), phytoplankton biomass (Selph *et al*., this issue), nutrients (NO_3_^−^ and NH_4_^+^, Knapp *et al*., this issue), and *in situ* incubation experiments were collected at 6 depths from the surface to the DCM on daily 02:00 a.m. (local U.S. central) CTD-Niskin rosette casts. Samples for shipboard incubations were collected from daily CTD casts near dusk. For more detail on sampling methodology see the aforementioned studies in this issue.

### Phytoplankton community composition and biomass

As reported in Selph *et al*. (this issue), we quantified phytoplankton abundance using a combination of flow cytometry (FCM), epifluorescence microscopy (EPI), and high-pressure liquid chromatography (HPLC). Cells identified by FCM were categorized into three populations (*Prochlorococcus* (PRO), *Synechococcus* (SYN), and picoeukaryotes (PEUK)) based on forward-angle and side-angle light scattering and fluorescence signatures for DNA, phycoerythrin, and chlorophyll. For larger cells (i.e., diatoms (DIAT), autotrophic dinoflagellates (ADINO), prymnesiophytes (PRYM)), and other eukaryotes (OTHER), a combination of HPLC and EPI was used to determine taxa-specific biomass. HPLC-derived pigments were partitioned into taxonomic groups using CHEMTAX (Wright *et al*., 2008; Higgins *et al*., 2011; Selph *et al*., this issue). EPI samples from the shallowest two depths (within the mixed-layer, which we define as the depth at which density increases by 0.125 kg/m^−3^ (Monterey and Levitus, 1997)) and the deepest two depths (just above and at the DCM) from three of the Lagrangian cycles (C1, C4, and C5) were used to determine depth−varying carbon:chlorophyll ratios, hence, carbon-based biomass for each group. Assuming Redfield C:N ratios (106:16, mol:mol), carbon-based biomasses from Selph *et al*., (this issue) were converted to nitrogen-based biomasses for the biogeochemical model (Redfield 1963). See Supplemental Appendix A and Selph *et al*. (this issue) for additional details.

### Phytoplankton growth, productivity, and grazing rates

We used three distinct incubation strategies for quantifying phytoplankton productivity and nutrient uptake rates: 24-h *in situ* incubations on the “incubation” array (NPP, NO_3_^−^ uptake, and taxon-specific phytoplankton growth rates); 6-h shipboard incubations starting at dawn (vertical patterns of NO_3_^−^ and NH_4_^+^ uptake); and diel shipboard incubations consisting of sequential 4-h incubations for 24 h (diurnal patterns of NO_3_^−^ and NH_4_^+^ uptake).

#### Taxon-specific growth rates

Two-point seawater dilution grazing experiments were conducted *in situ* daily at 6 depths spanning the euphotic zone to determine taxon-specific phytoplankton growth rates for a total of 88 independent experiments (Landry *et al*., 2008, 2011, this issue). 2.7-L samples of either whole seawater or partially diluted seawater (32% whole seawater/68% 0.1-μm filtered seawater) were incubated for 24-h on the incubation array. Initial and final samples for FCM (*Prochlorococcus* and *Synechococcus*), HPLC (dinoflagellates, diatoms and prymnesiophytes), and fluorometric chlorophyll *a* (bulk phytoplankton) were used to determine taxon-specific phytoplankton growth and mortality at each depth. See Supplementary Appendix A and Landry *et al*. (this issue) for additional details.

#### In situ NPP and nitrate uptake

NPP and ^15^NO_3_^−^ uptake rates were measured at the same 6 depths on the incubation array. Four incubation bottles (2.7-L) per depth were filled from Niskin rosettes (three light bottles, one dark bottle). All bottles were spiked with H^13^CO_3_^−^ (final concentration of 154 or 196 μmol L^−1^ on NF1704 and NF1802, respectively). Two light bottles were spiked with ^15^NO_3_^−^ (final concentration of 10 or 8 nmol L^−1^ on NF1704 and NF1802, respectively). Bottles were then incubated for 24 h on the array. Upon recovery, incubations were immediately vacuum filtered onto pre-combusted 25-mm GF/F filters in the dark. Filters were rinsed with filtered seawater, wrapped in foil and stored at −80°C. Samples were fumigated with HCl vapor to remove inorganic carbon, dried, and placed inside a tin cup to be used for C/N and isotopic analysis at the University of California, Davis stable isotope facility. Uptake rates (^15^NO_3_^−^ or bicarbonate) were determined for each incubation bottle using equations in Stukel (2020). NO_3_^−^ uptake is reported as the average and uncertainty of the ^15^NO_3_^−^ spiked bottles. As no statistically significant difference in H^13^CO_3_^−^ uptake was detected between bottles spiked with ^15^ NO_3_^−^ and those without, NPP is reported as the average and uncertainty of H^13^CO_3_^−^ uptake in the three light bottles corrected for dark bottle H^13^CO_3_^−^ uptake.

#### Shipboard vertically-resolved nitrate and ammonium uptake

Since NH_4_^+^ recycling within 24-h bottle incubations can substantially bias measurements of NH_4_^+^ uptake (Dugdale and Wilkerson, 1986), we conducted short-term (6-h) NH_4_^+^ and NO_3_^−^ uptake experiments in shipboard incubators. Each incubator was uniformly shaded using clear or blue-tinted acrylic sheets to achieve three light levels as determined by simultaneous measurements with a 2-π LI-COR photosynthetically active radiation (PAR) sensor for downwelling irradiance and a 4-π water-proof LI-COR PAR sensor for ambient irradiance in the incubator. These calibrations were done shipboard to account for light reflection or shading effects specific to their location. Incubation light levels for NF1704 (C1-C3) were determined to be 145% (clear, surface), 79% (mixed-layer), and 21% (lower mixed-layer) of surface irradiance (I_0_). For NF1802 (C4-C5), the clear incubator was replaced with one screened to 1.7% I_0_ to mimic DCM light. Incubators were cooled with mixed-layer seawater (24.5-26.5°C). Samples were drawn from depths near these light levels (as determined from noon CTD casts with CTD-mounted PAR sensor).

To determine patterns of nitrate and ammonium uptake as a function of depth, six 2.7-L samples from each of the three incubator light levels (e.g., 5, 15, and 45 m for surface, 79% and 21% I_0_, respectively) were collected near dusk and placed in the incubators until dawn (N.B., ship schedule and CTD water budget did not allow dawn sampling for these experiments). At dawn, triplicate samples from each depth were spiked with either ^15^NO_3_^−^ or ^15^NH_4_^+^ (concentrations as above) and incubated for 6-h (dawn to approximately local noon). Samples were then filtered and analyzed as described above.

#### Shipboard diel nitrate and ammonium uptake experiments

To determine diel variability in nutrient uptake and assess potential recycling occurring within 24-h incubations, we also conducted short-term (diurnally-resolved) nutrient uptake experiments. Nine to twelve 2.7-L bottles were filled at dusk. Two or three 24-h reference bottles (24-h incubation) and an additional “time point” bottle (4-h incubation) were immediately spiked with ^15^NO_3_^−^ or ^15^NH_4_^+^ (the remainder of the bottles were not immediately spiked). All bottles were then returned to the incubator. After ∼4 h, the first experimental bottle was removed from the incubator and filtered, and a second experimental bottle was spiked. This process continued for 24-h to produce six sequential 4-h incubations from which diurnal patterns of nutrient uptake could be determined. After 24-h, the reference bottles were harvested. No diel experiments were conducted during C3 (NF1704), and no diel ammonium uptake experiments were conducted during C4 (NF1802). On all other cycles (C1, C2, C5), simultaneous NO_3_^−^ or NH_4_^+^ uptake diel experiments were conducted. On NF1802 (C4-C5), diel experiment bottles were also spiked with H^13^CO_3_^−^ to simultaneously determine diel variability in NPP.

### Data assimilation and model parameterization

#### Model Structure

Phytoplankton growth is parameterized using transfer functions from the biogeochemical model NEMURO-GoM (Shropshire *et al*., 2020; Shropshire *et al*., this issue), which was designed for the open-ocean GoM. NEMURO-GoM parameterizes phytoplankton growth as a function of light, nutrients and temperature. By default, it includes two phytoplankton groups (small (SP) and large (LP)); however, we extended this model to parameterize 6 distinct groups that were identifiable through FCM, HPLC and/or EPI approaches above: (1) PRO, (2) SYN, (3) PRYM, (4) ADINO, (5) DIAT and (6) OTHER (equaling the residual biomass from the remaining groups of autotrophic eukaryotic phytoplankton).

NEMURO-GoM parameterizes group-specific growth rates with the following 7 equations: (1) NO_3_^−^ limitation (NL), (2) NH_4_^+^ limitation (AL), (3) light limitation (LL), (4) temperature limitation (TL), (5) respiration (Resp), (6) gross primary productivity (GPP), and (7) NPP, wherein K_NO3_ is the NO_3_^−^ half-saturation constant (µM), K_NH4_ is the NH_4_^+^ half-saturation constant (µM), α is the initial slope of the photosynthesis-vs-irradiance curve (m^2^ W^−1^ d^−1^), β is the light-inhibition constant (m^2^ W^−1^ d^−1^), and V is the maximum-specific-growth rate at a reference temperature (d^−1^). NEMURO-GoM uses a reference temperature of 0°C, but we provide values referenced to 25°C, which is representative of the GoM mixed-layer during our cruises. Q is the temperature effect coefficient (°C^−1^), R is biomass-specific respiration, E is growth-specific excretion, and B is biomass (mmol N m^−3^). Subscript “*i*” indicates group-specific parameters with nitrate concentrations (µM, [NO_3_^−^]), ammonium concentrations (µM, [NH_4_^+^]), PAR, and T (°C) as environmental conditions. For our Bayesian parameter-selection procedure (see below), all group-specific parameters except β were log transformed. Equations are as follows:

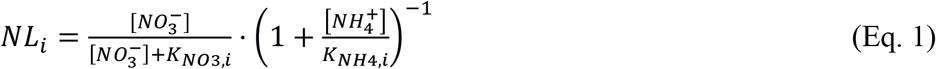

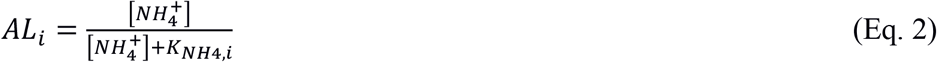

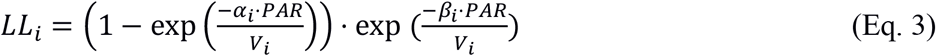

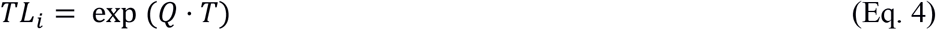

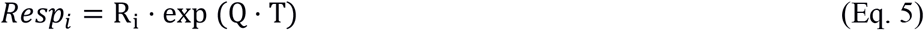

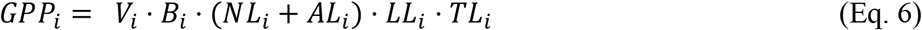

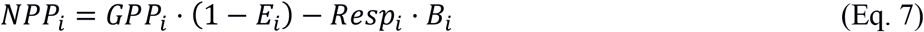

#### Bayesian Parameter Optimization Procedure

To constrain group-specific model parameters, we used a Bayesian parameter-selection method applied to a modified 0-dimensional version of NEMURO-GoM consisting of Eq. 1-7 with 6 phytoplankton taxa (additional details in Supplemental Appendix B). Our objective was not to find a single “best” parameterization, but rather to develop a statistical ensemble of possible parameter sets that reflects both uncertainty within the data and uncertainty in taxon-specific responses to light, temperature and nutrients. Briefly, the initial parameter set, *τ*_0_, is set to be equivalent to NEMURO-GoM default parameterizations for either SP or LP (Supplemental Table AI). The model is then initialized and run with *in situ* phytoplankton abundances, nutrient concentrations, temperature, and light equivalent to the observed initial conditions for every *in situ* incubation experiment. A joint probability is then used to assess both the model-data misfit (sum-of-squared normalized residuals) between observations (NPP, NO_3_^−^ uptake, and taxon-specific growth rates) and model results (Eq. 8; “likelihood”) and the prior probability of the parameter set used (Eq. 9).

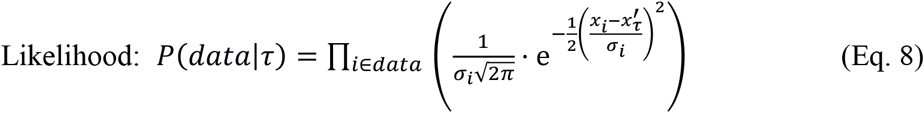

where x_*i*_ and σ_*i*_ are the measurement and measurement uncertainty, respectively, of the *i*’th observation and 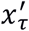 is the modeled value for that observation.

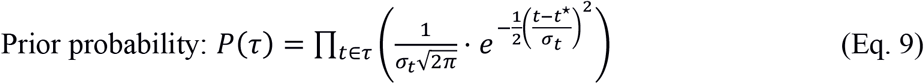

where *t*^*^ and σ_*t*_ are the mean and standard deviation, respectively, for the prior estimate of model parameter value *t* (Supplemental Table AI).

To explore the possible solution space, we used a Markov Chain Monte Carlo procedure with the Metropolis-Hastings algorithm (Hastings, 1970). For each iteration of the random walk, each parameter is varied by a small increment (in log-transformed space if a variable was transformed) yielding a new parameter set: *τ*_*j*+1_, which is used to re-run the model yielding a new likelihood and prior probability. The new parameter set would be “accepted” any time cost (i.e., −*P*(*data*|*τ*_*j*+1_)· *P*(*τ*_*j*+1_)) decreased and may be accepted even when cost increases based on the acceptance ratio: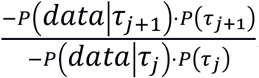 If the new parameter set is accepted, the random walk continues from *τ*_*j*+1_⋅ Otherwise a new *τ*_*j*+1_ is calculated. The first 50% of accepted parameter sets were removed (“burn-in”) and the remainder were subsampled (1 in 50) to remove autocorrelation between parameter sets and yielding ∼20,000 final parameter sets.

## RESULTS

### *In situ* conditions and the phytoplankton community

All five cycles had surface chlorophyll concentrations <0.2 µg L^−1^ and a pronounced DCM between 69 and 137 m (Girard *et al*., this issue). The top of the DCM typically corresponded to ∼1–2% I_0_. NO_3_^−^ concentrations were consistently <0.1 µM at depths shallower than 69 m and generally <1 µM throughout the entire euphotic zone (Knapp *et al*., this issue). Only C5, with the shallowest euphotic zone and closest to the shelf break, had >1 µM NO_3_^−^ at <100 m depth.

*Prochlorococcus* averaged ∼50% of the carbon-based phytoplankton biomass in all water parcels, although its proportional contribution varied from 31-66% of integrated euphotic zone biomass, being slightly more dominant in cycles with deeper DCMs (Table 2). PRO biomass typically increased with depth from the mixed-layer to a subsurface maximum slightly above the DCM before declining at the DCM. Mixed-layer SYN:PRO biomass ratios ranged from 0.2-1.4. However, SYN biomass decreased with depth, such that it represented ≤10% of PRO biomass at the DCM. Combined, cyanobacteria contributed a mean (± standard deviation) of 67±14% of euphotic zone integrated autotrophic carbon, while 2-10 μm eukaryotic cells comprised most of the remaining biomass. Prymnesiophytes were the dominant eukaryotic taxa, comprising 17 ± 7%, while chlorophytes and pelagophytes represented 5±3% and 4.2±2%, respectively, of integrated euphotic zone carbon biomass. Diatoms were only minor contributors to autotrophic biomass (<2% in all cycles). For additional details on community composition, see Selph *et al*. (this issue).

**Table I.** Summary of data for each Lagrangian cycle. Surface NO_3_^−^ and NH_4_^+^ are averaged across days within a cycle. NPP, NO3− and *f*-ratio are vertically integrated values. *f*-ratio = NO_3_^−^ uptake / (NH_4_^+^ uptake + NO_3_^−^ uptake) is a blended product that combines shipboard NO_3_^−^ and ammonium uptake experiments (more accurate) with *in situ* NO_3_^−^ uptake and NPP measurements (greater depth resolution).

|  | Cycle 1 | Cycle 2 | Cycle 3 | Cycle 4 | Cycle 5 |
| --- | --- | --- | --- | --- | --- |
| **Surface $\text{NO}_3^-$**<br>**($\mu\text{M}$)** | <0.1 | <0.1 | <0.1 | <0.1 | <0.1 |
| **Surface $\text{NH}_4^+$**<br>**($\mu\text{M}$)** | <0.025 | <0.025 | <0.025 | <0.025 | 0.025 |
| **Surface Chl *a***<br>**($\mu\text{g/L}$)** | 0.107±0.04 | 0.049 | 0.080±0.007 | 0.100±0.01 | 0.127±0.01 |
| **NPP**<br>**($\text{mmol C m}^{-2} \text{d}^{-1}$)** | 25.6±0.1 | 24.4±2.8 | 24.9±0.4 | 29.2±3.0 | 30.1± 2.9 |
| ***In Situ* $\text{NO}_3^-$**<br>**Uptake**<br>**($\mu\text{mol N m}^{-2} \text{d}^{-1}$)** | 3.45±1.63 | 1.49±0.21 | 0.99±0.37 | 0.74±0.02 | 1.48±0.09 |
| ***f*-ratio** | 0.22 | 0.02 | 0.05 | 0.06 | 0.12 |

**Table II.** Summary of *in situ* observations. Group-specific biomass are vertically integrated to the base of the euphotic zone for each Lagrangian cycle. Units are mmol N m^−2^ (converted from carbon-based values reported in Selph *et al*. (this issue) by dividing by a Redfield C:N ratio of 106:16, mol:mol). Values marked with NM were not measured.

|  | Cycle 1 | Cycle 2 | Cycle 3 | Cycle 4 | Cycle 5 |
| --- | --- | --- | --- | --- | --- |
| *Prochlorococcus* | 6.16 ± 0.22 | 6.76 ± 0.28 | 6.85 ± 0.37 | 4.85 ± 0.04 | 3.24 ± 0.45 |
| *Synechococcus* | 1.44 ± 0.04 | 1.25 ± 0.03 | 1.22 ± 0.04 | 2.57 ± 0.05 | 2.36 ± 0.37 |
| **Diatom** | 0.26 ± 0.11 | NM | NM | 0.07 ± 0.004 | 0.04 ± 0.003 |
| **Dinoflagellate** | 0.47 ± 0.05 | NM | NM | 0.40 ± 0.11 | 0.43 ± 0.02 |
| **Prymnesiophytes** | 0.33 ± 0.03 | NM | NM | 0.44 ± 0.18 | 0.54 ± 0.03 |
| **Other Eukaryotes** | 2.46 ± 0.84 | 0.79 ± 0.0 | 0.60 ± 0.0 | 0.47 ± 0.11 | 1.64 ± 0.5 |

### Vertical profiles of NPP and nutrient uptake

NPP declined almost monotonically with depth on all cycles, although mixed-layer NPP was typically only 2 to 3-fold higher than NPP at the DCM (Fig. 1). NPP showed relatively weak inter-cycle variability. Cycle-average NPP varied from 0.27-0.55 μmol C L^−1^ d^−1^ at 5 m and varied from 0.10-0.29 μmol C L^−1^ d^−1^ at the DCM. The highest vertically integrated NPP (mean: 29.3 mmol C m^−2^ d^−1^) was for C5 (the cycle closest to the shelf), which also had the shallowest DCM, euphotic zone depth, and nitracline. The lowest vertically−integrated NPP was measured during C2 (24.3 mmol C m^−2^ d^−1^).

**Figure 1.**
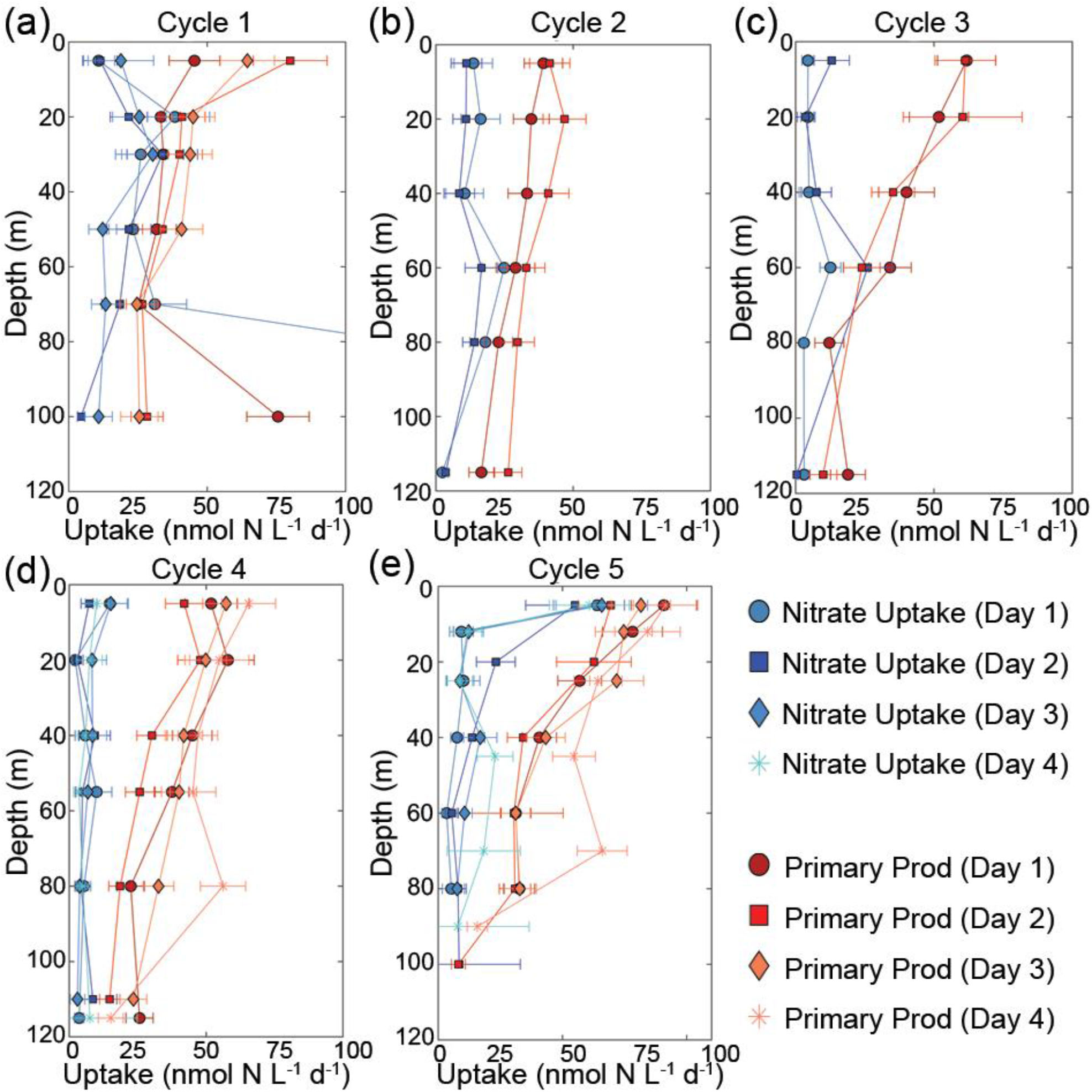
NO_3_^−^ (red) and HCO_3_^−^ uptake (blue) rates from *in situ* incubations. HCO_3_^−^ uptake rates are converted to nitrogen units assuming Redfield ratio (6.625 molar ratios of C:N). Error bars represent 1 standard deviation of the mean.

Vertical structure was less distinct for NO_3_^−^ uptake rates (Fig. 1). Across depths and cycles, NO_3_^−^ uptake typically ranged from 5−20 nmol N L^−1^ d^−1^. C1 exhibited a modest subsurface peak at 20−30 m (the rate was much higher for one outlier experiment at the DCM, likely due to contamination). C2 and C3 showed evidence for a weak subsurface maximum at ∼60 m, while rates were relatively constant with depth for C4. C5 was the only cycle with a clear surface maximum in NO_3_^−^ uptake rates. Vertically integrated, cycle−average NO_3_^−^ uptake ranged from 0.72 (C4) to 3.45 mmol N m^−2^ d^−1^ (C1).

To investigate the relative proportion of NPP supported by NO_3_^−^ uptake (commonly referred to as the *f*-ratio), we computed an *f*-ratio from *in situ* data as:

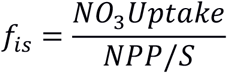

where S is Redfield C:N stoichiometry (106:16). We caution that while *f*_*is*_ is similar to the commonly used *f*-ratio (proportion of N uptake provided by NO_3_^−^), it should not be interpreted as equivalent to the technical definition of the *f*-ratio (new production divided by total production), because we suspect that nitrification in the euphotic zone may have been significant (Kelly *et al*., in review; Stukel *et al*. this issue b). Furthermore, *f*_*is*_ may overestimate an *f*-ratio calculated from NO_3_^−^ and NH_4_^+^ uptake (see below), if phytoplankton engage in luxury nutrient uptake. With those caveats, vertically-integrated, cycle-average *f*_*is*_ ranged from 0.17-0.89; however, the high value (0.89 from C1), was heavily biased by the aforementioned high NO_3_^−^ uptake outlier at the DCM. If the results from this day’s experiments are excluded, *f*_*is*_ for this cycle decreases to 0.47. Across all cycles, *f*_*is*_ generally increased with depth in the euphotic zone. It is important to note that heterotrophic bacteria may play a role in nutrient uptake; however, the data suggests that ammonium uptake was vertically-correlated with net primary production and phytoplankton growth rates (i.e., ammonium uptake was low at the light-limited DCM), while nitrate uptake was negligible during the night. Since community nitrate and ammonium uptake were light limited, we suspect that the majority of nutrient uptake was mediated by phytoplankton.

Shipboard, depth-resolved NO_3_^−^ and NH_4_^+^ uptake experiments showed that NH_4_^+^ uptake is consistently higher than NO_3_^−^ uptake for all cycles and depths (Fig. 2). Similar to the findings from *in situ* experiments, the data suggests that NH_4_^+^ is the preferred nitrogen source. Experiments during 2017 (mixed-layer only) showed little variability with depth, although NO_3_^−^ uptake was higher in C1 than C2 (as suggested by *f*_*is*_). NO_3_^−^ and NH_4_^+^ uptake rates from 2018 (which extended to the depth of the DCM) showed that NH_4_^+^ uptake was lower at the DCM than in the mixed-layer, a finding that also agrees with our *in situ* NPP results. Shipboard incubation-based *f*-ratios (*f*_*deck*_ = NO_3_^−^uptake / (NO_3_^−^uptake + NH_4_^+^uptake)) were lower than *f*_*is*_ and ranged from 0.02-0.17. For f_deck_, we assumed that NH_4_^+^ uptake occurs at constant rates over the full 24-h daily period (and hence multiplied 6-h experiments by 4), but that NO_3_^−^ uptake occurs only during the day (hence multiplied 6-h experiments by 2), based on the results of diel experiments (next section). Because *f*_*deck*_ calculated from NO_3_^−^ and NH_4_^+^ uptake measurements should be considered more accurate due to the shorter incubation times with less recycling occurring, we developed a blended *f*-ratio product: we treated *f*_*deck*_ as more accurate at paired depths, but used the vertical patterns from *f*_*is*_. That is, we multiplied *f*_*is*_ by the ratio of *f*_*deck*_:*f*_*is*_ at the nearest depths. This approach suggested that euphotic zone-averaged *f*-ratios varied from 0.02-0.22 (mean=0.06). *f*-ratios were slightly higher in the deep euphotic zone (>50 m depth, *f* ranged from 0.03-0.44, mean=0.14) than in the shallow euphotic zone (*f* = 0.01-0.14, mean=0.06).

**Figure 2.**
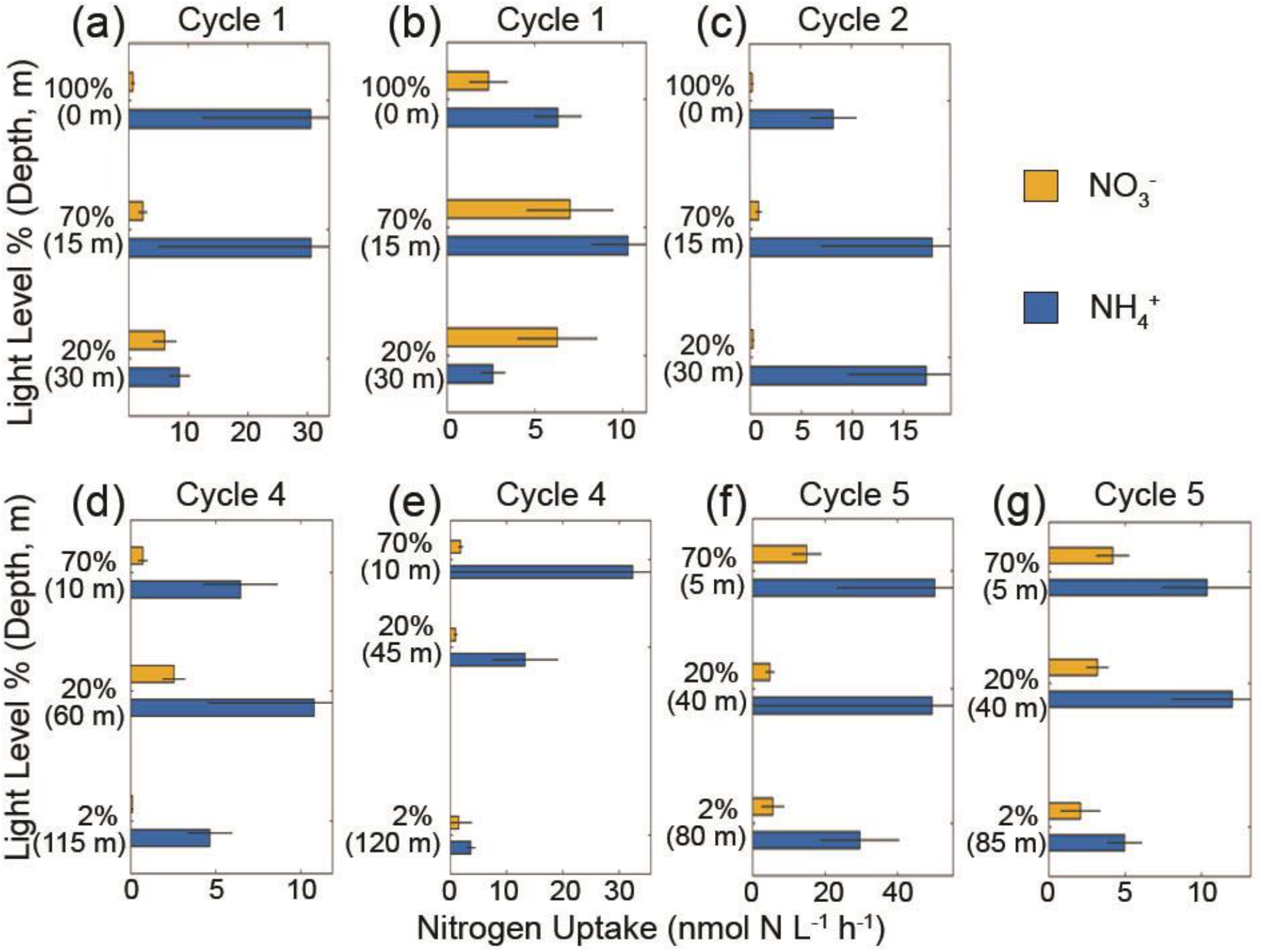
Results from NO_3_^−^ (blue) and NH_4_^+^ (orange) uptake experiments conducted shipboard at three light levels and water collection depth (m) as indicated. Error bars are replicate measurements.

### Diel variability in NPP and nutrient uptake

Distinct patterns emerged from the sequential 4-h incubations assessing diel variability in NH_4_^+^ uptake (Fig. 3), NO_3_^−^ uptake (Fig. 4), and NPP (Fig. 5). NO_3_^−^ uptake and NPP were strongly light-dependent, with consistent mid-day peaks and low activity during the night (Figs. 4, 5). However, NH_4_^+^ uptake showed no distinct diel patterns, with uptake rates as high at night as during the day. Comparison to simultaneous 24-h incubations allowed us to assess potential nutrient recycling in the euphotic zone. For NPP, the sum of six sequential 4-h H^13^CO_3_^−^ uptake experiments were generally not significantly different from the mean of simultaneous 24-h uptake experiments, indicating, as expected, that the time scale of H^13^CO_3_^−^ recycling within the euphotic zone is long relative to the duration of a 24-h experiment. However, for both NO_3_^−^ uptake and NH_4_^+^ uptake rates, the sums of 4-h incubations were substantially greater than simultaneous 24-h incubations. For NO_3_^−^ uptake, the ratio of the sum of sequential 4-h incubations to the average of 24-h incubations ranged from 1.2-5.6 (median=2.0). For NH_4_^+^ uptake this ratio ranged from 5.0-6.0 (median=5.7). These results indicate rapid recycling of NH_4_^+^ in the euphotic zone, and also reasonably fast recycling of NO_3_^−^.

**Figure 3.**
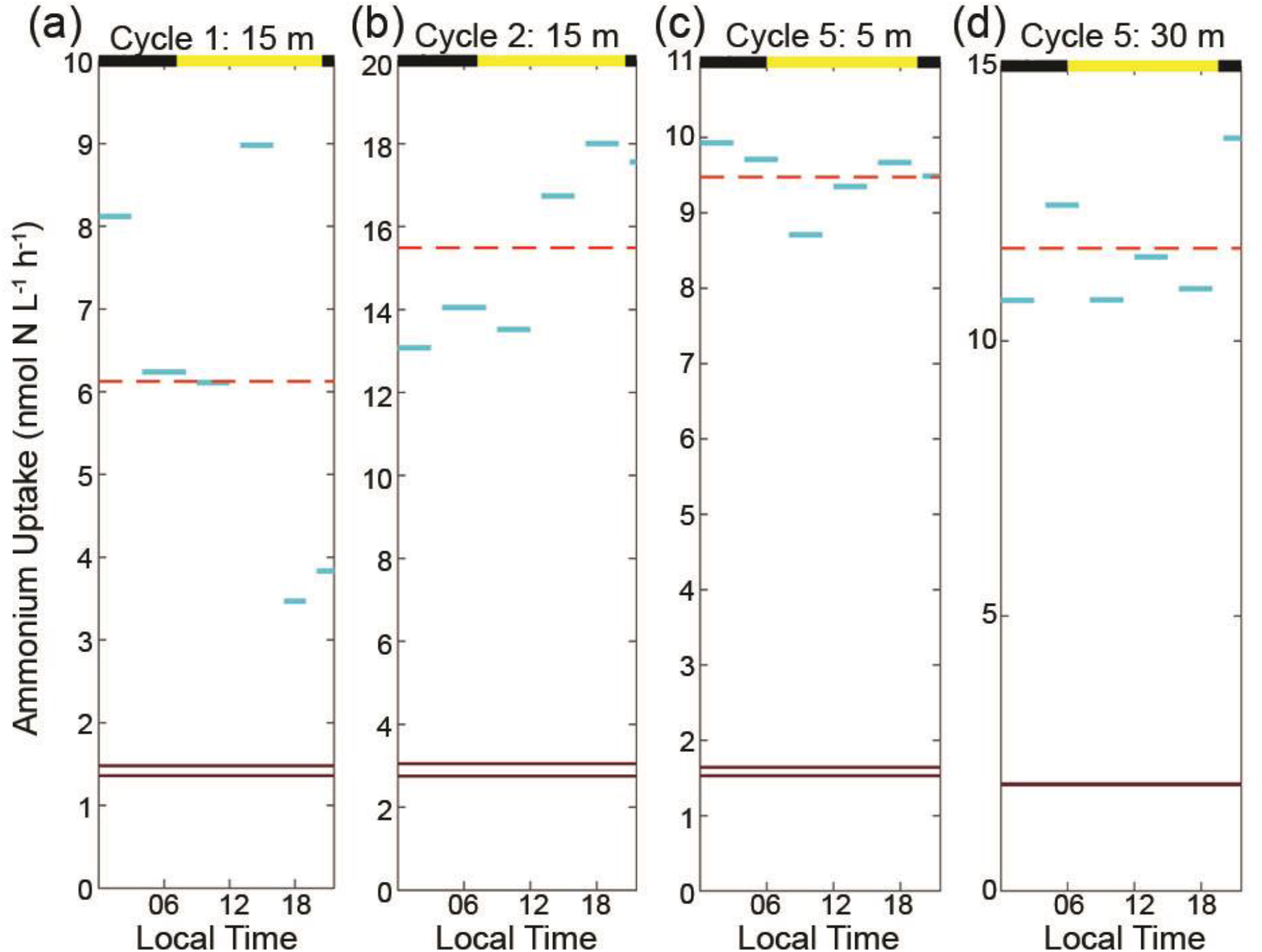
Diel NH_4_^+^ uptake rates (nmol N L^−1^ h^−1^) with solid horizontal lines showing the duration and mean uptake rates for the 4-5 h (blue) and 24-h (dark brown) incubations. Dashed horizontal lines (orange) represent averages of 4-5-h incubations for comparison with 24-h incubations carried out during the same time frame. Black and yellow alternating bars (top of each figure) represent local daylight and dark hours. Depths indicate depth of water collection and approximate correspondence to incubator light level.

**Figure 4.**
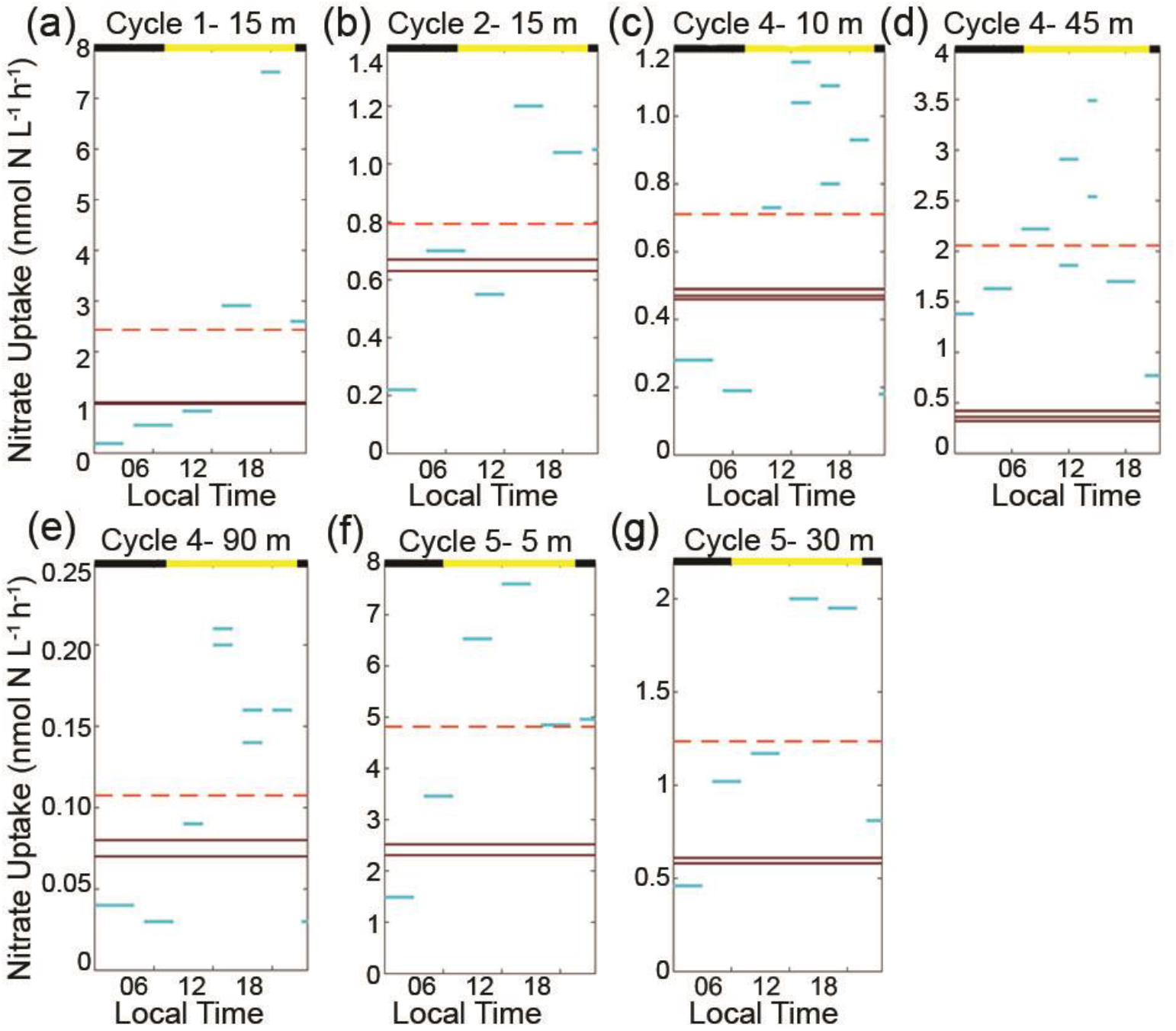
Diel NO_3_^−^ uptake rates (nmol N L^−1^ h^−1^). Depths of water collection for the incubations are indicated next to the cycle number. Solid horizontal lines show the duration and mean uptake rates for the 4-5-h (blue) and 24-h (dark brown) incubations. Dashed horizontal lines (orange) represent averages of 4-5-h incubations for comparison with 24-h incubations carried out during the same time frame. Time periods that had more than one replicate are both shown, e.g, two blue lines at one time period. Black and yellow alternating bars (top of each figure) represent local daylight and dark hours.

**Figure 5.**
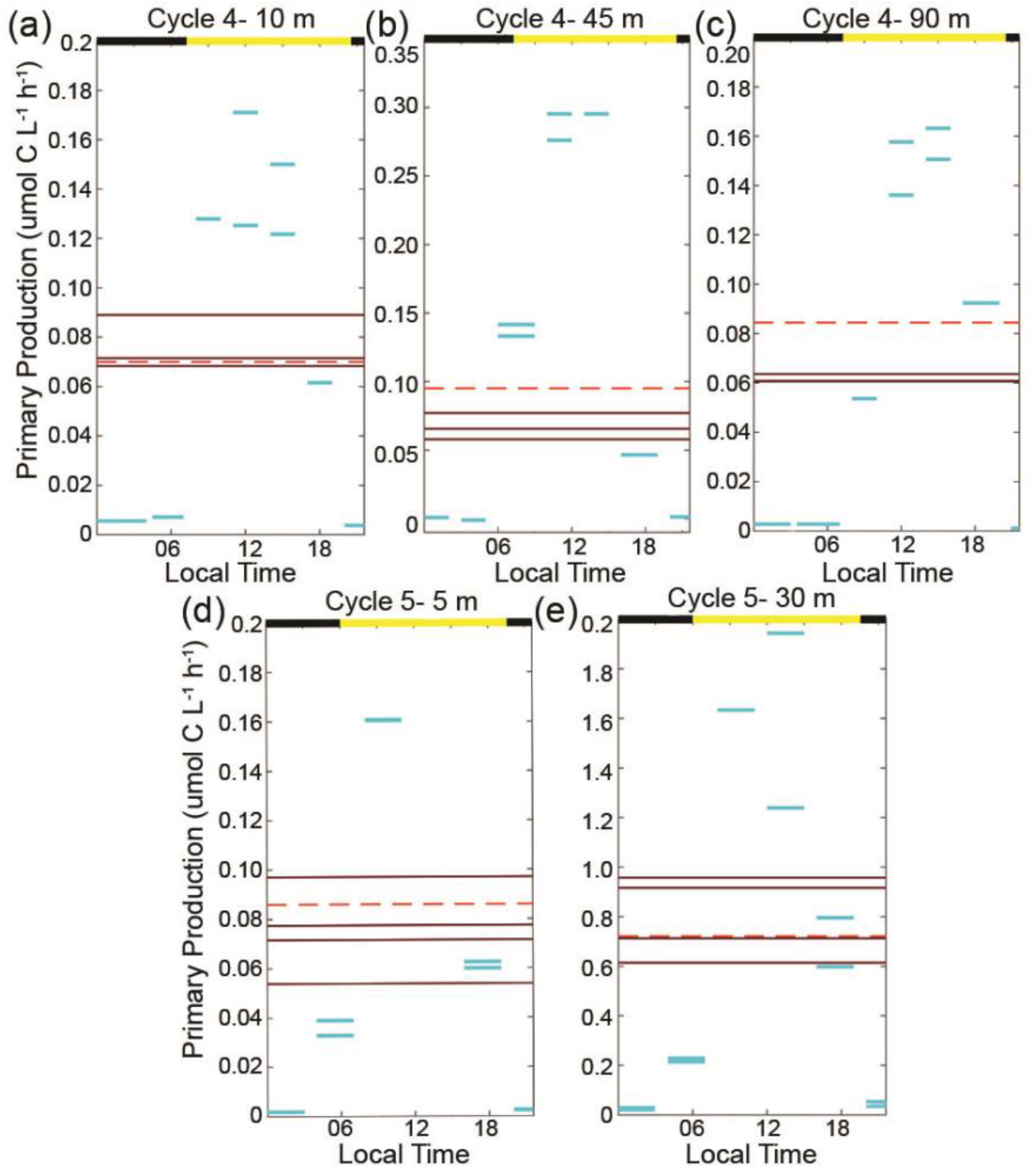
Diel primary production (µmol C L^−1^ h^−1^) from shipboard experiments. Solid horizontal lines show the duration and uptake rates for individual 4-5 h (blue) and duplicate 24-h (dark brown) incubations. Time periods that had more than one replicate are both shown, e.g, two blue lines at one time period. Dashed horizontal lines (orange) represent averaged short incubations for comparison with 24-hour incubations. Black and yellow bars located at the top of figures represents local daytime and nighttime.

### Vertical profiles of phytoplankton growth rates

Instantaneous growth rates of phytoplankton were variable as a function of depth and cycle (Fig. 6). On average, PRO surface growth rates were 0.67±0.35 d^−1^, decreasing with depth to 0.27±0.10 d^−1^ at the DCM. Similarly, the average SYN surface growth rates were 0.64±0.29 d^−1^, decreasing with depth to 0.34±0.30 d^−1^ at the DCM. These rates are consistent with previous research (Partensky *et al*., 1999). DIAT growth, ranging from −0.4 to 1.5 d^−1^, was higher in the upper euphotic zone and declined gradually with depth, except for a peak at 85 m during C4. ADINO growth ranged from 0-1.2 d^−1^ and varied between cruises. In 2017 (C1-C3), growth of ADINOs was highest in the upper mixed-layer and gradually declined with depth. In 2018 experiments, (C4-C5), ADINO growth peaked in the mid euphotic zone at ∼40 m. PRYM growth rate varied among cycles, ranging from −0.5 to 1.3 d^−1^ with occasional peak values in the mid water column.

**Figure 6.**
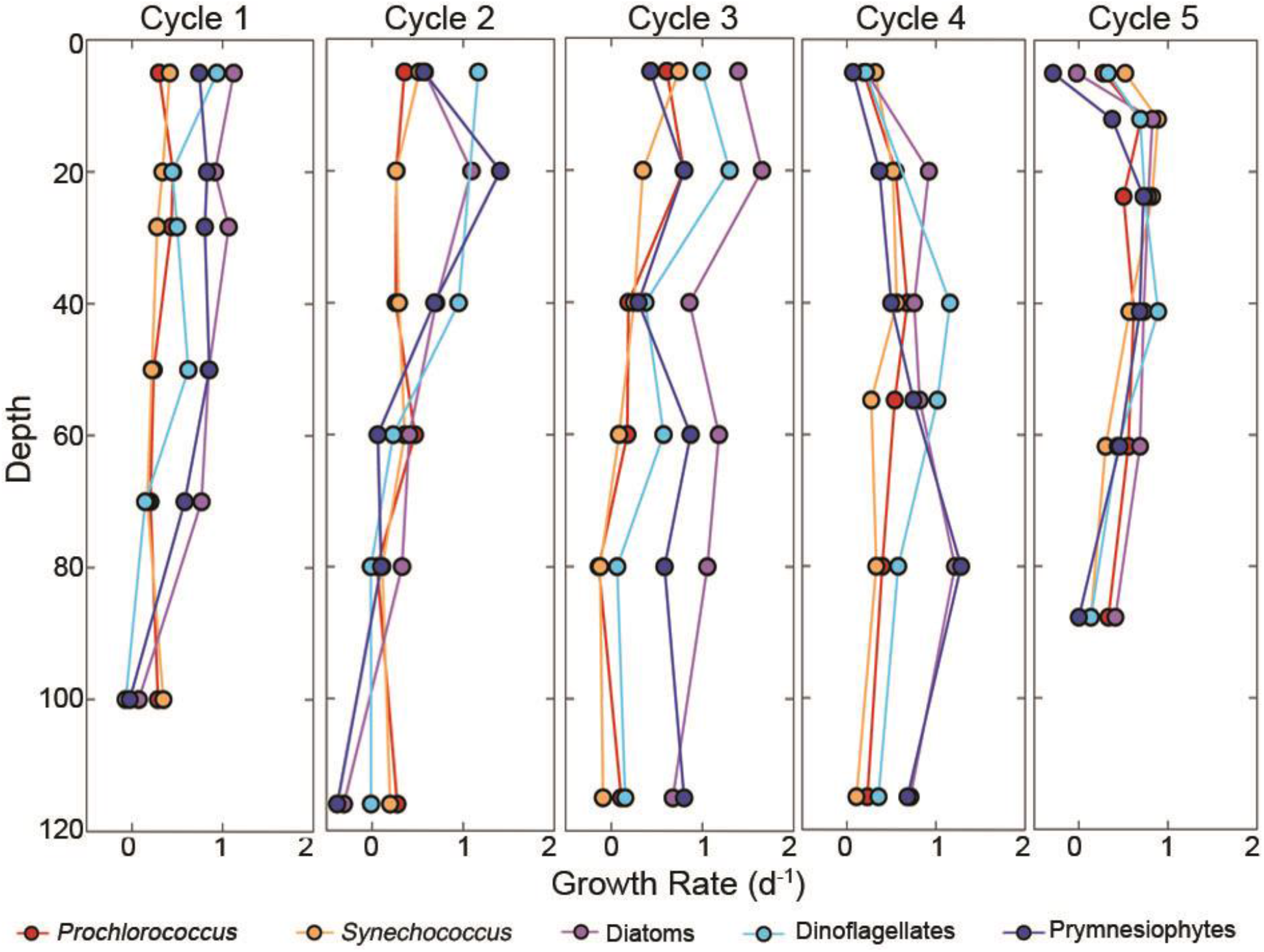
Dilution experiment results from cycles 1-5, showing averaged group-specific growth rates for *Prochlorococcus, Synechococcus*, diatoms, dinoflagellates and prymnesiophytes as a function of depth. Details of experiments are in Landry *et al*., this issue.

### Model results

Using a Bayesian statistical approach to assimilate our *in situ* measurements, we constrained typical transfer functions used to model taxon−specific phytoplankton responses to light, temperature, and nutrient limitation in biogeochemical models. Data assimilation increased the fit to experimental data relative to the default parameterization from NEMURO-GoM (Supp. Fig. 2; Supp. Table I). The parameters determined from data assimilation had a much better fit to NPP observations than the default parameters, but only a moderately better fit to nitrate uptake observations (Fig. 7). The assimilated parameters underestimated observed nitrate uptake when nitrate uptake was high (although only by a factor of 2–5, while the default parameters underestimated nitrate uptake by ∼10-fold). The model underestimate resulted from the fact that even with low half-saturation constants, the NEMURO model did not predict nitrate uptake rates as high as the observations in a region with such low nitrate concentrations. Total model log-likelihood increased substantially after the Bayesian optimization procedure (from −40,748 to −3,333 for NEMURO-GOM default and data-assimilated parameters, respectively, Supp. Table I). Notably, the default parameters predicted unrealistically low NPP and growth rates for phytoplankton based on the low nutrient concentrations in the GoM euphotic zone (Supp. Fig. 2). Consequently, the optimized parameter ranges suggested much lower K_NH4_ (NH_4_^+^ half-saturation constants) for all phytoplankton taxa except SYN (Fig. 8). K_NH4_ was lowest for PRO (mean = 0.007 µM, 95% C.I. = 0.005–0.009 µM), which thrives in oligotrophic regions. Previous research supports low half-saturation values for PRO (Marañón *et al*., 2013; Grossowicz *et al*., 2017). In contrast, the posterior distributions for K_NO3_ remained relatively close to the prior distributions (and to default NEMURO-GoM parameterizations, Fig. 8). The Bayesian parameter optimization approach suggested K_NO3_ values of ∼3 μmol L^−1^ for all large phytoplankton taxa, ∼0.4 μmol L^−1^ for PRO and OTHER, and ∼0.1 μmol L^−1^ for SYN. This is comparable to laboratory studies that suggest half saturation constants are proportional to cell size with larger cells ranging in value from 0.1−5.0 μmol L^−1^ (Eppley *et al*., 1969).

**Figure 7.**
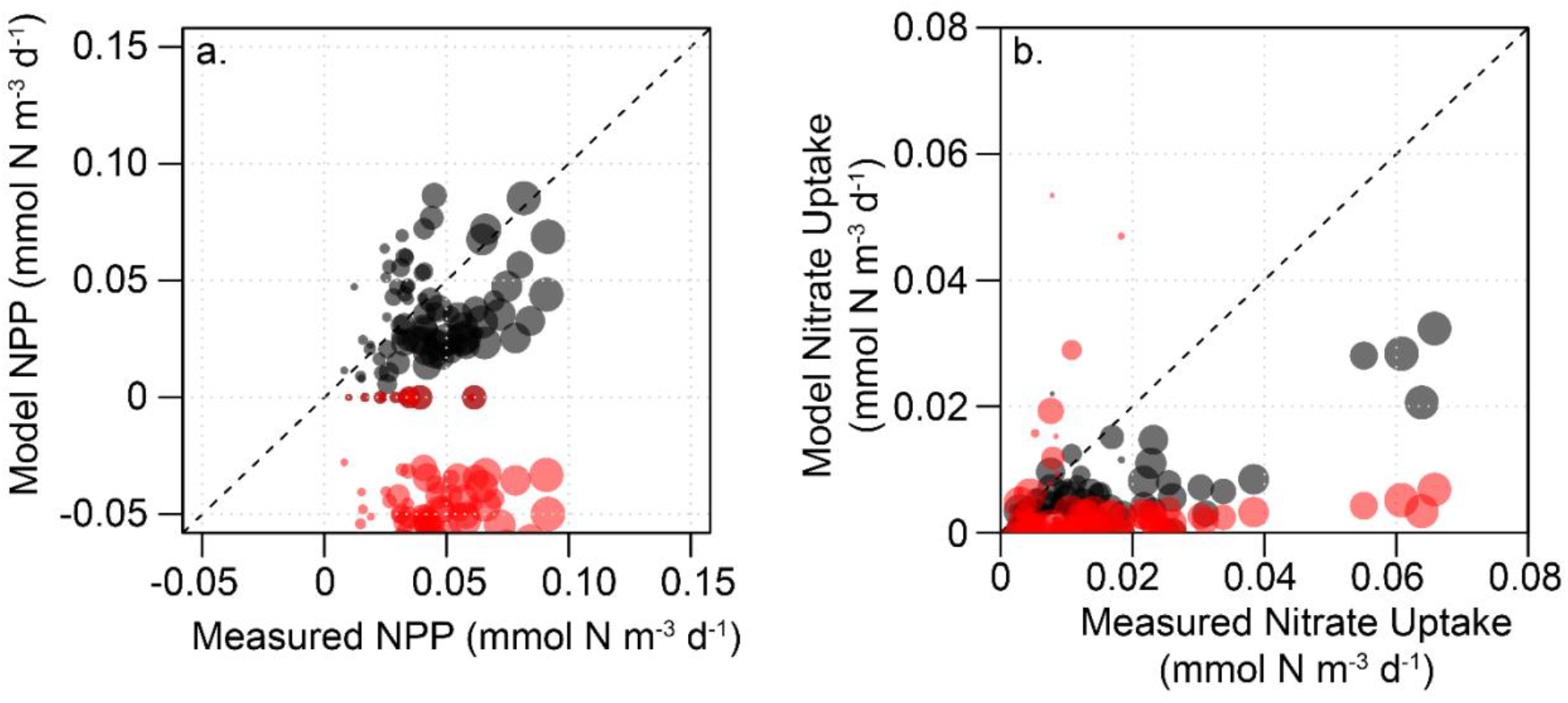
Model-observation comparisons for (a) NPP and (b) NO_3_^−^ uptake. The dashed line represents the 1:1 ratio. Red dots indicate NEMURO-GoM parameterization while black dots show the “optimized” parameterization.

**Figure 8.**
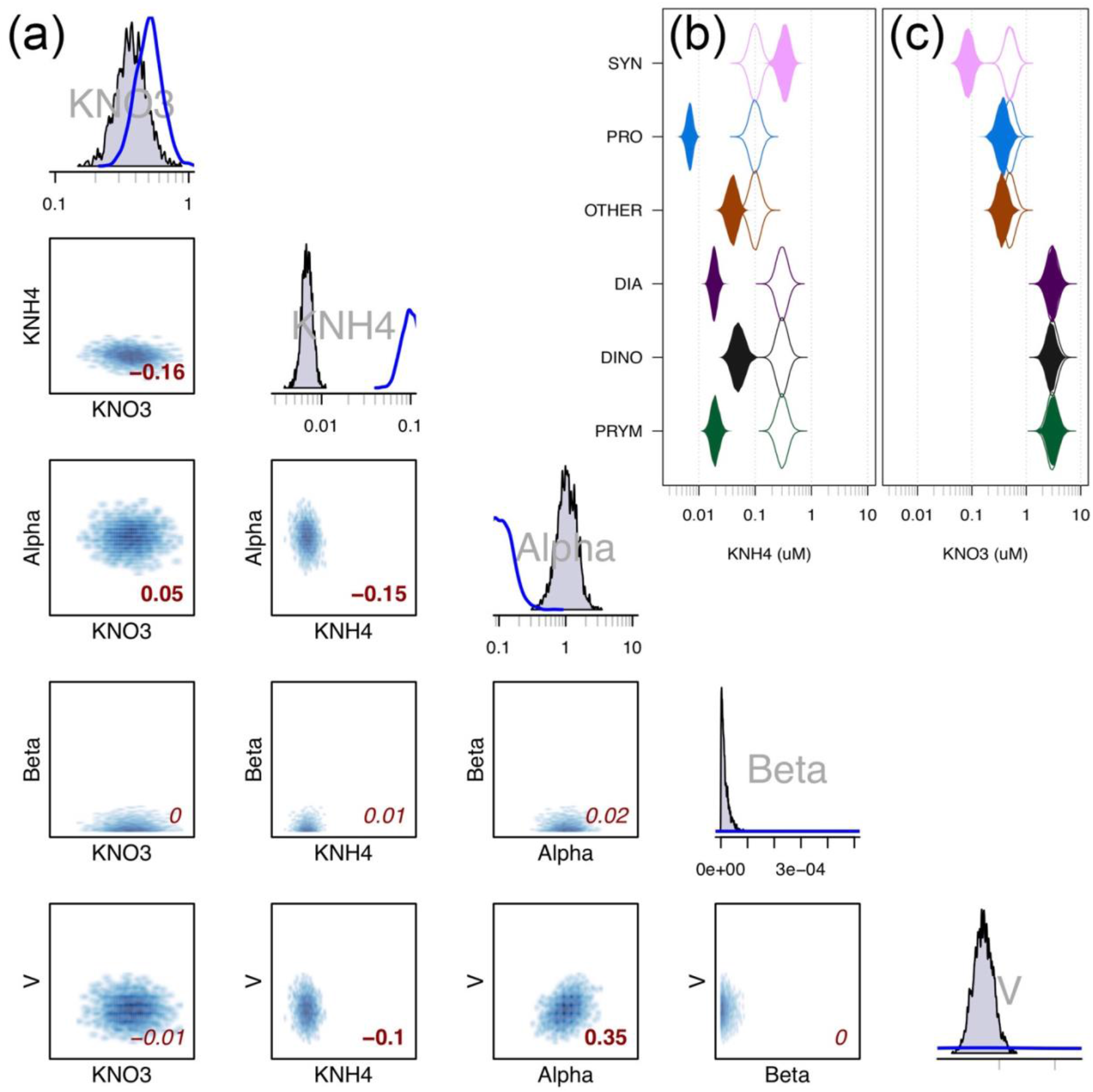
(a) Model parameters of log-transformed K_NH4_ (μmol L^−1^), log-transformed K_NO3_ (μmol L^−1^), log-transformed Alpha (m^2^ W^−1^ d^−1^), Beta (m^2^ W^−1^ d^−1^) and log-transformed V at 0°C (d^−1^) for *Prochlorococcus*. Upper plots are histograms for each variable (filled gray lines, n = 10^6^, subsampled to 2×10^4^), with blue lines approximating the prior distribution that was assumed for each variable from previous studies. Lower property-property plots show the correlation between any two parameters (note that scales are as in the histograms above). Red value located in bottom right corner in each subplot gives Pearson’s correlation coefficient (which is suitable for use with non-normal data, Nefzger and Drazow, 1957), with bold values indicating significance at p < 0.05. Violin plots of **(b)** NH_4_^+^ half-saturation constants (K_NH4_, µM) and **(c)** NO_3_^−^ half-saturation constants (K_NO3_, µM) from the optimized model (filled) and their priors (empty).

Model results also varied among groups with respect to light-response parameters and maximum growth rates. PRO and OTHER had the lowest maximum growth rates at 25°C (V_25°C_ = 0.82 d^−1^, 95% C.I.=0.73-0.92 d^−1^ and 0.44 d^−1^, 95% C.I.=0.35-0.54 d^−1^, respectively as shown in Supplementary Fig. 2). However, the low maximum growth rate parameter suggested for OTHER might be an artifact of the inclusion of many different taxa within this group, which resulted in the model predicting low and only weakly-varying growth rates. All large phytoplankton taxa had a maximum growth rate at 25°C of approximately 1.6-2.0 d^−1^, while SYN had a higher maximum growth rate of 2.6-4.4 d^−1^. PRO had the weakest light sensitivity, with α (initial slope of the photosynthesis-irradiance relationship) equal to 0.01 m^2^ W^−1^ d^−1^ (95%: 0.006−0.02 m^2^ W^−1^ d^−1^), which likely reflects the presence of low-light-adapted PRO strains that thrive in the deep euphotic zone. Most taxa had α in the range of 0.02-0.8 m^2^ W^−1^ d^−1^ reflecting substantial light limitation and reduced growth rates in the DCM (Fig. 8, Supp. Fig. 1).

Because we simultaneously varied all parameters, our optimization procedure provides some insight into correlations among parameters. A subset of these results (for PRO) are shown in Fig. 8. For instance, α and V were positively correlated suggesting that the model could fit PRO growth rates at the DCM with either a higher maximum growth rate or with a weaker sensitivity to light limitation. Correlations for PRO were relatively weak for most parameter pairs. SYN, however, shows strong correlations for several parameter pairs. In particular, nutrient half-saturation constants were positively correlated with each other and with V. This suggests that the model could determine viable solution sets with high maximum growth rates and substantial nutrient limitation or with low maximum growth rates and weaker nutrient limitation, and that the relative importance of nutrient limitation by NO_3_^−^ or NH_4_^+^ could vary (although K_NO3_ was always lower than K_NH4_ for SYN).

Comparison of optimized parameter sets to measured environmental conditions (nutrients, light, and temperature) also allowed us to assess patterns of limiting resources for each taxon (Fig. 9). While all phytoplankton taxa experienced slightly greater nutrient limitation in the shallow euphotic zone than in the deep euphotic zone, with the exception of SYN, nutrient limitation led to <50% decrease in growth rates (Fig. 9a). SYN was the only group with a median *f*-ratio greater than 0.05. This suggests that all taxa (except SYN) rely disproportionately on ammonium as a N source for growth and primary productivity. In contrast, SYN appeared to be the sole NO_3_^−^ specialist, with commensurately high *f*-ratios (∼0.5) and greater nutrient limitation of growth rates. Our model thus shows a distinct niche specialization of the abundant taxa to the oligotrophic nature of this ecosystem.

**Figure 9.**
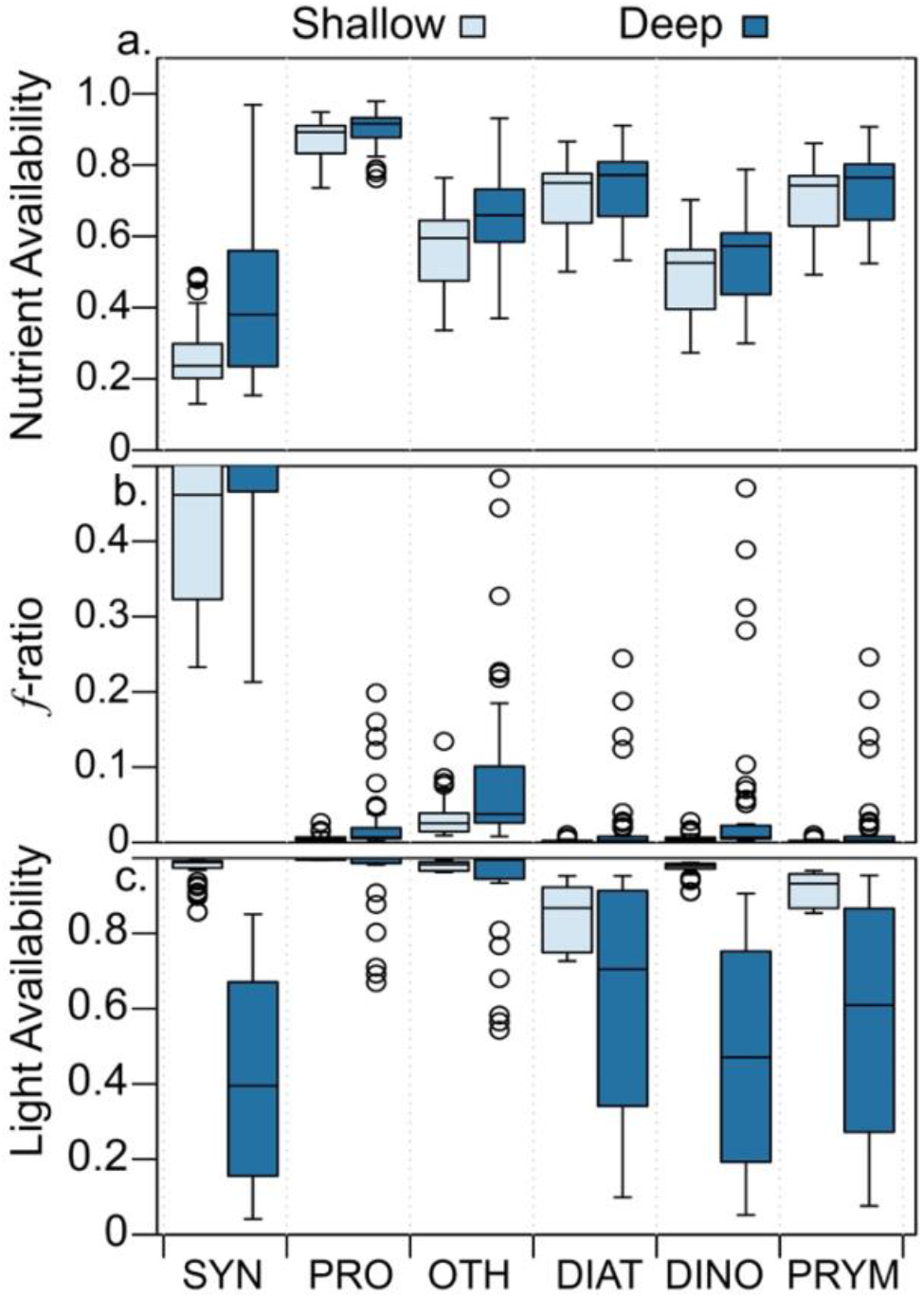
Modeled results for each phytoplankton group for (a) nutrient availability (unitless growth rate multiplier equal to NL + AL in Eqs. 1 and 2), (b) *f*-ratio and (c) light availability (unitless growth rate multiplier equal to LL in Eq.) as a function of depth. Light bars represent shallow water (< 50 m) and dark bars represent deeper samples (>50 m). Note that the upper quartile and 95% C.I. values are offscale for SYN in panel b with values of 0.551 and 0.696 for shallow and 0.699 and 0.921 for deep. See equations 1-4 for formulas and additional information on the calculations.

With the exception of PRO and OTHER, which seem to be well-adapted to maintaining low and comparatively insensitive growth rates in both the mixed-layer and the deep euphotic zone, all taxa experienced substantial light limitation in the deep euphotic zone (>50 m). At these depths, light limitation ranged from mild limitation at ∼50 m depth (e.g., growth penalties of ∼20%) to strong limitation at the DCM (growth penalties of up to 95%). In comparison, no groups exhibited substantial light-limitation above 50 m; the greatest average light-limitation penalty in the upper 50 m was for DIAT (∼10%), which had the highest photo-inhibition parameter.

## DISCUSSION

### Diel and vertical variability in phytoplankton productivity and nutrient utilization

Our data indicates that the oligotrophic GoM is a highly-stratified, picophytoplankton-dominated (mostly PRO) region, with low nitrate concentrations (typically <0.1 μM above the DCM), low chlorophyll *a* concentrations (typically <0.03 μg L^−1^ in the mixed-layer and <0.2 μg L^−1^ in the DCM), and deep DCMs (69 to 137 m). Nutrient uptake measurements show that NH_4_^+^ recycling is key to sustaining the phytoplankton community and productivity. These conclusions are consistent with previous investigations of phytoplankton nutrient limitation in the oceanic GoM (Dugdale and Goering, 1967; Eppley and Peterson, 1979; Platt and Harrison, 1985; Lipschultz, 2001; Wawrik *et al*., 2004).

Vertical profiles of NPP (Fig. 1) and taxon−specific growth rates (Fig. 6) indicate that phytoplankton and nutrient uptake are generally light limited, with NPP and growth rates declining with depth. This decrease in NPP was not associated with a decline in phytoplankton biomass. Although the biomass of different taxa varied with depth, overall phytoplankton biomass and POC were typically relatively constant or increased slightly with depth (Selph *et al*., this issue; Stukel *et al*., this issue a). This decrease in productivity and growth is more gradual, however, than might be expected based on light limitation alone. Instead, it reflects substantial nutrient limitation in surface waters that gradually gives way to light limitation in the deeper euphotic zone, suggesting that phytoplankton throughout the euphotic zone grow at rates substantially below their physiological potential. Vertical trends also likely include shifts in species composition. Although these patterns are not evident from our FCM and HPLC-based taxonomic study, sequencing data has shown differentially adapted species within our broad categories, especially for high-light or low-light adapted PRO and SYN (Gutiérrez-Rodríguez *et al*., 2016). Notably, the distinct DCM was primarily, but not solely, driven by increased cellular pigmentation, as has been seen in multiple other studies (Cullen *et al*., 1982; Mignot *et al*., 2014). The maintenance of the DCM despite reduced growth rates thus requires either reduced mortality (i.e., grazing) or subsidies from sinking phytoplankton (Hodges and Rudnick, 2004; Moeller *et al*., 2019; Stukel *et al*., this issue a).

The diel experiments showed distinct temporal patterns of NH_4_^+^ and NO_3_^−^ uptake. NH_4_^+^ uptake rates did not vary with time of day (Fig. 3), while NO_3_^−^ uptake rates were light-dependent and similar to diel patterns of NPP (Figs. 4 and 5). This is consistent with NO_3_^−^ uptake being more coupled to photosynthetic energy generation while NH_4_^+^ uptake had no cellular regulation (Figs. 3 and 4) (Dortch, 1990). Similar patterns have been found for NO_3_^−^ uptake in the Costa Rica Dome (Stukel *et al*., 2016). However, NO_3_^−^ uptake rates measured in the Sargasso Sea were lower at night but remained at ∼1/3 daytime values (Lipschultz *et al*., 2001). In the Sargasso Sea, different studies have also given different ratios of daytime to nighttime NH_4_^+^ uptake, with Lipschultz *et al*. (2001) determining that daytime rates were twice those of nighttime rates, while Glibert *et al*. (1988) found negligible day-night differences (as in our study). Lipschultz *et al*. (2001) methods included longer incubations (dusk to dawn) while Glibert *et al*. (1988) included short 1-2 h incubations. Taken together, these results underline the difficulty in extrapolating from short (2-6 h) experiments to daily uptake rates or *f*-ratios. Our diel uptake experiments also showed substantial differences between 4-h and 24-h NH_4_^+^ uptake incubations (and to a lesser, but still considerable, extent for NO_3_^−^ uptake incubations). Rapid cycling of NH_4_^+^ in the surface ocean is expected, while NO_3_^−^ is often considered upwelled “new” nitrogen (Dugdale and Goering, 1967). Recent evidence, however, suggests that NO_3_^−^ regeneration through NH_4_^+^ oxidation may be more common in surface oceans than previously realized (Yool *et al*., 2007, Shiozaki *et al*., 2016). The combination of relatively high rates of NO_3_^−^ uptake (compared to NO_3_^−^ concentration), very low NO_3_^−^ inventory from the surface to the DCM at ∼100 m, and the ∼2-fold higher NO_3_^−^ uptake measured in 4-h incubations relative to 24-h incubations lead us to believe that NO_3_^−^ in the surface GoM might be supplied primarily by NH_4_^+^ oxidation (Stukel *et al*., this issue b).

### Model Optimization

Accurately constraining biogeochemical model parameters is a challenging yet crucial step towards treating models as falsifiable hypotheses (Arhonditsis and Brett, 2004; Anderson, 2005; Franks, 2009; Schartau *et al*., 2017). Objective parameterization typically follows two broad approaches: (1) empirical fits of specific transfer functions to available data, or, (2) formal data assimilation. The former approach typically fits simple functional forms (e.g., photosynthesis-irradiance curves or Ivlev grazing formulations) to field or laboratory data. Empirical fits can be developed using experimental data derived from intentional manipulation of independent variables for a specific species or community (Eppley *et al*., 1969; Platt *et al*., 1980), or determined by fitting relationships for measurements made under natural conditions across an environmental gradient (Li *et al*., 2010; Morrow *et al*., 2018). These two empirical approaches address subtly different questions. The former approach examines the mechanistic response of a single taxon to changes in an environmental variable, while the latter addresses a similar question but at a community-level scale, thus accounting for community shifts that occur with changes in light and nutrient conditions. This investigation of mechanistic responses at the community level is often more appropriate for models, because most such models simulate plankton functional groups comprised of many different species (Hood *et al*., 2006). However, it is complicated by frequent covariance of multiple parameters (e.g., light and nutrients).

The alternative approach to model parameterization, which has long been common in physical oceanography, is formal data assimilation (Friedrichs *et al*., 2006; Gregg *et al*., 2009; Cummings *et al*., 2013). In data assimilation, the full forward model is typically run and compared to observations, and then a prescriptive approach is followed to sequentially modify a subset of the parameters to minimize the model-data misfit. While multiple approaches have been used (Lawson *et al*., 1995; Ward *et al*., 2010; Doron *et al*., 2011), the most common approach is the variational-adjoint, which is frequently used to compare model outputs to spatial maps of sea surface chlorophyll or time-series measurements of standing stocks in a one-dimensional framework. However, some studies have questioned whether assimilation of standing stock measurements is sufficient to constrain many model parameters, or have noted that more complex models often respond less well to assimilation techniques and the tendency of the variational-adjoint method to gravitate toward local minima in parameter space (Xiao and Friedrichs, 2014; Löptien and Dietze, 2015).

Our Bayesian statistical approach allows us to simultaneously constrain multiple parameters for which *a priori* estimates contained substantial uncertainty. The incorporation of multiple types of rate measurements allows for more precise determination of posterior distributions than would have been possible if we only utilized standing stock measurements. Indeed, optimized parameter values for the NEMURO-GoM transfer functions suggest >90% reduction in cost on average (Supp. Table I). The results returned reasonable parameter value ranges for most taxa, and demonstrated some expected emergent properties (e.g., PRO is an oligotrophic specialist). They also, however, disagreed with *a priori* expectations in some key ways, indicating that likelihood (i.e., our measurements) was more important than prior distributions in determining the posterior distributions. Most notably, the model showed that our experimental results could only be matched using much lower NH_4_^+^ half-saturation values for most taxa than the default values in NEMURO-GoM or common half-saturation constants measured in cultures (Edwards *et al*., 2012; Shropshire *et al*., 2020).

Nevertheless, some parameters seem unrealistic. Notably, the OTHER group, which is a composite of multiple phytoplankton taxa including pelagophytes, chlorophytes, prasinophytes, cryptophytes and other taxa, had very low maximum growth rates (mean: 0.44 d^−1^, 95%: 0.35 – 0.54 d^−1^ at 25°C) and very little nutrient or light limitation. This suggests that the optimization procedure attempted to give this composite group a low and relatively constant growth rate under all conditions. It is unlikely that pelagophytes, chlorophytes, prasinophytes, and cryptophytes were all insensitive to environmental variability. More likely, the different taxa within OTHERS have different responses to light and nutrients that are masked when they are aggregated together in a single group. On the other hand, this aggregate behavior might also be considered a strength of this parameterization approach, since even the most accurate representation of any one taxon or group can never reflect the adaptive physiological potential of a natural mixed community. It is also important to consider that groups like ADINO and OTHERS might be mixotrophic and taking up nitrogen by phagotrophy in this region, which may impact their functional responses to light and nutrient limitation (Jones 2000; Stoecker *et al*., 2017).

Even though posterior distributions were very different from prior distributions, many of the parameters (e.g., K_NH4_) retained substantial uncertainties. These uncertainties arise, in part, from covariance among environmental variables, which leads to correlations across the different parameters. In future work, experimental manipulations could be conducted in ways that break some of the natural correlations (e.g., nutrient amendments or light manipulation). In addition, some uncertainty undoubtedly arises from errors in measurements (nutrients, light, phytoplankton abundance). Future work could explicitly formulate the model such that the inputs are “true” values of standing stocks, defined as 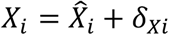 for incubation *i*, where 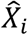 is the measured value and *δ*_*Xi*_ is a random variable with prior distribution centered at 0 and a standard deviation equal to the measurement uncertainty. The *δ*_*Xi*_ would then be variables to solve for parameter solutions. Even without these potential improvements, however, we believe that our approach provides useful guidance for parameterizing future biogeochemical models of the GoM and elsewhere.

### Phytoplankton niches in the oligotrophic ocean

The vast number of species, strains, or ecotypes that are aggregated into our broad phytoplankton taxonomic groups make it difficult to define specific niches for each group (Scanlan, 2003; Simon *et al*., 2009; Kashtan *et al*., 2014). It is reasonable to expect that the evolutionary history of taxa provides some inherited physiological constraints on their overall niche space, while the prevalence of specific ecotypes may contribute further to a definable niche space for each group. Our results, with respect to both spatial patterns of abundance and model constrained characteristics, shed light on these questions.

The two cyanobacterial taxa have distinctly different ecological niches. For PRO, the consistent subsurface maxima just above the DCM (Selph *et al*., this issue) and high α (initial slope of the photosynthesis-irradiance curve) indicates that it thrives in low-light conditions. We also observed a substantial increase in divinyl chlorophyll *a* cell^−1^ with depth that likely explains its physiological adaptation to low light (whether by cellular regulation or distinct ecotypes remains unconstrained). Other studies have shown that there are distinct depth distributions of PRO ecotypes (HLI and HLII near the surface, and LL at low light) (Zwirglmaier *et al*., 2007). Beyond light limitation, the very low NH_4_^+^ half−saturation constant shows it to be a low-nutrient specialist that relies almost exclusively on rapidly recycled NH_4_^+^. This concurs with a general consensus that although some strains of PRO contain genes for NO_3_^−^ assimilation (Martiny *et al*., 2009), this genus primarily utilizes NH_4_^+^ (Moore *et al*., 2002). In contrast, SYN was the only group that efficiently exploited low nitrate concentrations, typically maintaining a shallower distribution in the water column with greater sensitivity to light limitation. We thus suspect that SYN relies substantially on NO_3_^−^ regenerated in the upper euphotic zone through shallow nitrification (Kelly *et al*., submitted; Stukel *et al*., this issue b). Previous research has shown similar fundamental differences between SYN and PRO in nutrient utilization, with SYN utilizing nitrate more efficiently than PRO (Scanlan and West, 2002). Comparisons of abundances to nutrient concentrations and turbulent mixing rates have also suggested that PRO is more abundant in warm waters with low nitrate supply, while SYN and picoeukaryote niches are in cooler waters with greater nitrate supply (Otero-Ferrer *et al*., 2018), a finding that agrees with previous evidence showing PRO dominance in oligotrophic gyres and SYN and picoeukaryotes becoming more important in equatorial upwelling and temperate regions (Zubkov *et al*., 2000; Landry and Kirchman, 2002).

Model results for eukaryotic phytoplankton taxa were slightly less conclusive. The larger uncertainty is partially due to the substantially lower abundance of these taxa, and also in part because the pigment-based growth rates were imperfect estimates of true cellular growth of these taxa. DIAT had modest maximum growth rates, lower than those in many culture and field measurements (e.g., Furnas, 1990; Sarthou *et al*., 2005; Selph *et al*., 2011; Selph *et al*., 2016). This might, however, reflect chronic extreme silicon limitation, which was not assessed in our study. We do note that prior concentration measurements from GoM indicate that Si is very low (Dortch and Whitledge, 1992). Irwin *et al*. (2012) used continuous-plankton-recorder-derived phytoplankton abundance data to show that diatoms and dinoflagellates have distinct ecological niches with diatoms typically excelling in cold, nutrient-rich, well-mixed, low-light environments relative to dinoflagellates, which agrees with the low diatom biomasses we found. Barton *et al*. (2015) further showed that diatom responses to these physical drivers are spatiotemporally variable, which does suggest caution when applying our results to other regions. Our model could not significantly differentiate between ADINO and PRYMN, both of which had maximum growth rates of ∼0.6 d ^−1^ and exhibited substantial light limitation. The model did, however, suggest that the NH_4_^+^ half-saturation constant was substantially lower for PRYMN than ADINO, which is not surprising for a generally smaller phytoplankton taxon and might help explain why PRYM was the biomass-dominant eukaryote in our study (Selph *et al*., this issue). Considered together, these results delineate distinct competitive differences between these diverse phytoplankton functional groups, which helps explain their coexistence in oligotrophic conditions and supports the advanced light and nutrient resource competition model (Burson et al 2018).

## CONCLUSION

The oceanic GoM is an extremely oligotrophic ecosystem with low NPP and a strong DCM. We found higher NPP in the upper euphotic zone than the DCM, fueled mostly by recycled nutrients. Ammonium uptake exceeded nitrate uptake at all depths and was relatively invariant with time of day, while nitrate uptake was mainly restricted to daytime. Bayesian parameter optimization techniques allowed us to constrain maximum growth rates, light utilization parameters, and half saturation constants for five phytoplankton taxa (PRO, SYN, DIAT, PRYMN and ADINO), yielding parameter values for future modeling studies. This approach also allowed us to define distinct niches for different taxa and determine that all, except PRO, were chronically nutrient limited at all depths.

## ACKNOWLEDGEMENTS

This manuscript would not have been possible without the support from the captain, crew and research technicians aboard the NOAA Ship *Nancy Foster*; thank you for your dedication and hard work. In addition, we would like to acknowledge and thank our collaborators at NOAA Southeast Fisheries/RASMAS/University of Miami for their effort and perseverance on leading two successful cruises.

## DATA ARCHIVING

Data presented here have been submitted to the National Oceanic and Atmospheric Administration’s (NOAA) National Centers for Environmental Information (NCEI) data repository and will also be archived at BCO-DMO (Biological and Chemical Oceanography Data Management Office) site https://www.bco-dmo.org/program/819631. Model code and dataset are freely available on Github (https://github.com/tbrycekelly/Yingling/) with instructions on how to replicate model results.

