## Supplementary figures 1 and 2 and Supplementary Information for "Taxon-specific phytoplankton growth, nutrient utilization, and light limitation in the oligotrophic Gulf of Mexico"

### Supplemental Materials

#### Supplemental Appendix A. Methods for phytoplankton taxon-specific biomass and growth rates

Taxon-specific phytoplankton carbon biomass was determined using a combination of flow cytometry (FCM), high-pressure liquid chromatography (HPLC), and epifluorescence microscopy (EPI). Daily samples (2-mL) were collected from 6 depths spanning the euphotic zone, preserved with 0.5% paraformaldehyde, and frozen in liquid $N_2$ for post-cruise FCM analysis (Selph *et al.*, this issue). Samples were thawed and stained with DNA stain Hoechst 33342 and analyzed on a Beckman Coulter EPICS Altra flow cytometer (Monger and Landry, 1993; Selph *et al.*, 2011). Cells were categorized into three populations (*Prochlorococcus* (PRO), *Synechococcus* (SYN), and picoeukaryotes (PEUK)) based on fluorescence signatures for DNA, phycoerythrin, and chlorophyll, as well as forward- and side-scatter. PRO and SYN were assumed to have a carbon content of 32 and 101 fg C cell$^{-1}$ (Garrison *et al.* 2000; Brown *et al.* 2008)

Larger cells (diatoms (DIAT), autotrophic dinoflagellates (ADINO), prymnesiophytes (PRYM), and other eukaryotes (OTHER)) were quantified using a combination of HPLC and EPI as described in Selph *et al.* (this issue). HPLC samples (2.2-L) were collected from the same 6 depths as FCM samples, filtered onto GF/F filters, and frozen in liquid $N_2$. Samples were analyzed at the Horn Point Analytical Services Laboratory. Resulting pigments (monovinyl chl a, divinyl chl a, monovinyl chl b, divinyl chl b, chl c3, zeaxanthin, fucoxanthin, 19'hex-fucoxanthin, 19'but-fucoxanthin, allophycocyanin, peridinin, neoxanthin, and prasinoxanthin) were partitioned into taxonomic groups using CHEMTAX (Wright *et al.*, 2008; Higgins *et al.*, 2011; Selph *et al.*, this issue), providing chlorophyll-based estimates of the relative contributions of each of these groups. We then used EPI to determine carbon:chlorophyll ratios and carbon-based biomasses for these groups. EPI samples (450 mL filtered through 8-µm filters for microphytoplankton analysis, 50 mL filtered through 0.8-µm filters for nanophytoplankton analysis) from the shallowest two depths (mixed-layer) and the deepest two depths (just above and at the DCM) from multiple casts on three of the Lagrangian cycles (C1, C4, and C5) were stained with DAPI (protein stain) and proflavin (protein stain). Phytoplankton cells were determined based on DAPI-fluorescence and chlorophyll autofluorescence and cell outlines were visualized using proflavin fluorescence, allowing estimates of cellular dimensions for biovolume determinations. Biovolume was converted to cell carbon content using carbon:volume conversion equations (Menden-Deuer and Lessard, 2000). Eukaryotic phytoplankton biomass was then divided by the total eukaryotic contribution to chl a (from CHEMTAX, above) to determine depth-varying eukaryotic carbon:chl ratios. These ratios were multiplied by the respective contribution of each phytoplankton group (DIA, ADINO, PRYM, OTHER) to determine that group's carbon biomass at each depth. Assuming Redfield ratio (106:16, C:N), carbon concentrations were converted to mmol N $m^{-2}$ in order to have consistency with units for the biogeochemical model (Redfield 1963). For additional details, see Selph *et al.* (this issue).

To determine *in situ* growth rates of the same phytoplankton groups (SYN, PRO, DIA, ADINO, PRYM, OTHER) we conducted two-point microzooplankton grazing pressure dilution experiments (Landry *et al.* 1982; Landry *et al.* 1984; Landry *et al.* 2008). Seawater samples were collected from the same six depths as biomass measurements described above. Initial samples were analyzed for FCM and HPLC as described above. One "control" bottle (2.7-L) was filled unamended with natural seawater. A second "treatment" bottle was filled with 32% natural seawater and 68% 0.1-μm filtered seawater from the same depth. Both samples were placed in mesh bags and incubated at the depth from which the sample was taken on a drifting *in situ* array to ensure natural temperature and light conditions (Landry *et al.*, 2009). After 24 h, the array was recovered and samples were filtered for FCM and HPLC "finals" from each control and treatment bottle. Taxon-specific phytoplankton growth rates were then determined from the differences in taxon-specific net growth rates in control and treatment bottles. These experiments were conducted each day of every cycle. For additional details, see Landry *et al.* (this issue).

#### Supplemental Appendix B. Methods for Bayesian Optimization Procedure

In general, the posterior probability is proportional (to some constant) to the likelihood and prior probability (Eq. A1). This fact allows for estimation of the former as long as the latter can be calculated. While several sampling schemes exist, we implemented the Metropolis-Hastings algorithm, which uses a Markov Chain Monte Carlo random walk to sample the solution space.

$$P(\tau|data) \propto P(data|\tau) \cdot P(\tau) \qquad \text{(Eq. A1)}$$

Starting with a parameter set defined by NEMURO-GOM, $\tau_0$, the likelihood (i.e., $P(data|\tau)$; Eq. 8) and prior probability (i.e., $P(\tau)$; Eq. 9) are calculated. A candidate parameter set is generated by adding a small value to each parameter with values sampled from a normal distribution centered on zero and a standard deviation chosen for each parameter based on the relative sensitivity of the model to that parameter (e.g., the model was sensitive to β so a small step size was used (1 x $10^{-6}$) compared to $K_{NO3}$ of 0.01). The likelihood and prior probability for the candidate parameter set, $\tau_{i+1}$, are calculated. The acceptance ratio, $\alpha$, for the candidate parameter set is given in Eq. A2:

$$\alpha = \min\left(1, \frac{-P(data|\tau_{i+1}) \cdot P(\tau_{i+1})}{-P(data|\tau_i) \cdot P(\tau_i)}\right) \qquad \text{(Eq. A2)}$$

If the candidate parameter set is accepted, the random walk continues from $\tau_{i+1}$, producing a new candidate parameter set. If it is not accepted, a new candidate is formed from $\tau_i$. Note that the average acceptance ratio is sensitive to step size and so was used to initially set the standard deviation used above (to yield an acceptance ratio of ~0.1 – 0.4).

It was important to choose priors for each parameter that reflected the distribution of values observed in marine phytoplankton. To do this, we first log-transformed appropriate parameters (Supplemental Table AI) in order to work with normally distributed parameters and values wherever possible. We used one set of priors for DIAT, PRYM, ADINO and a separate set of priors for PRO, SYN, OTHER (Supplemental Table AI). All values were estimated from Edwards et al. (2012) except for α, which was estimated from Bouman *et al.* (2018), and β, which was given a uniform ($\beta \in U(0{,}0.01)$ $m^2$ $W^{-1}$ $d^{-1}$) prior due to the large sensitivity of the model to changes in light inhibition.

**Supplemental Table AI.** Priors used for each parameter in the model given as either a normal distribution (N(μ, σ)) or as a uniform distribution (U(min,max)). Parameters which were log-transformed are prefixed by $\log_{10}$. Groups are *Prochlorococcus* (PRO), *Synechococcus* (SYN), other autotrophic eukaryotes (OTHER), diatoms (DIAT), autotrophic dinoflagellates (ADINO), and prymnesiophytes (PRYM).

| **Parameter** | **PRO** | **SYN** | **OTHER** | **DIAT** | **ADINO** | **PRYM** |
|---|---|---|---|---|---|---|
| $\log_{10}$KNO3 | N(-0.3, 0.1) | N(-0.3, 0.1) | N(-0.3, 0.1) | N(0.48, 0.1) | N(0.48, 0.1) | N(0.48, 0.1) |
| $\log_{10}$KNH4 | N(-1, 0.1) | N(-1, 0.1) | N(-1, 0.1) | N(-0.52, 0.1) | N(-0.52, 0.1) | N(-0.52, 0.1) |
| $\log_{10}$Alpha | N(-1, 0.2) | N(-1, 0.2) | N(-1, 0.2) | N(-1, 0.2) | N(-1, 0.2) | N(-1, 0.2) |
| Beta | U(0, 0.01) | U(0, 0.01) | U(0, 0.01) | U(0, 0.01) | U(0, 0.01) | U(0, 0.01) |
| $\log_{10}$V | N(-0.4, 0.5) | N(-0.4, 0.5) | N(-0.40, 0.5) | N(-0.10, 0.5) | N(-0.10, 0.5) | N(-0.10, 0.5) |
| $\log_{10}$R | N(-1.05, 0.2) | N(-1.05, 0.2) | N(-1.05, 0.2) | N(-1.52, 0.2) | N(-1.52, 0.2) | N(-1.52, 0.2) |

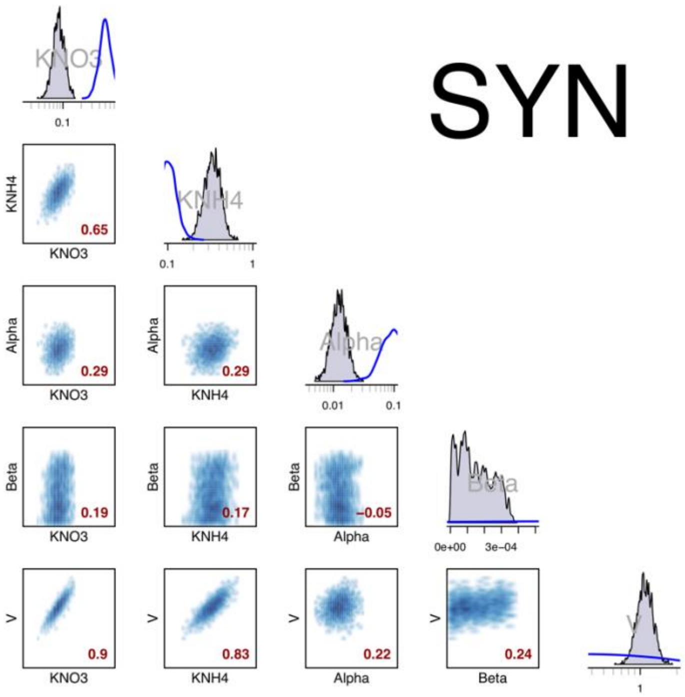
SYN
KNO3
KNH4
Alpha
Beta
V
0.65
0.29
0.29
0.19
0.17
−0.05
0.9
0.83
0.22
0.24
0.1
1
0.01
0e+00
3e−04

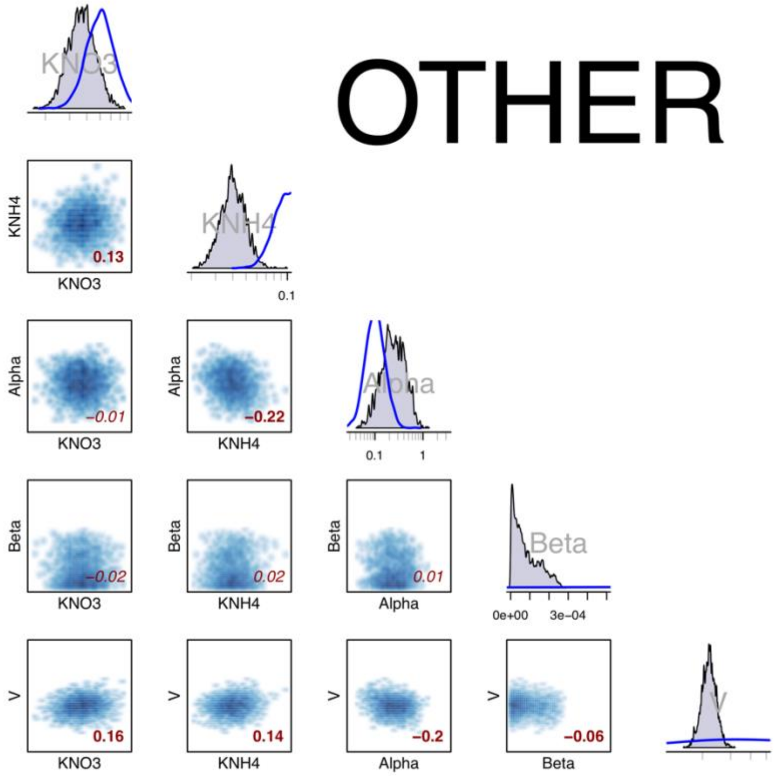
OTHER
KNO3
KNH4
Alpha
Beta
V
0.13
−0.01
−0.22
−0.02
0.02
0.01
0.16
0.14
−0.2
−0.06
0.1
1
0e+00
3e−04

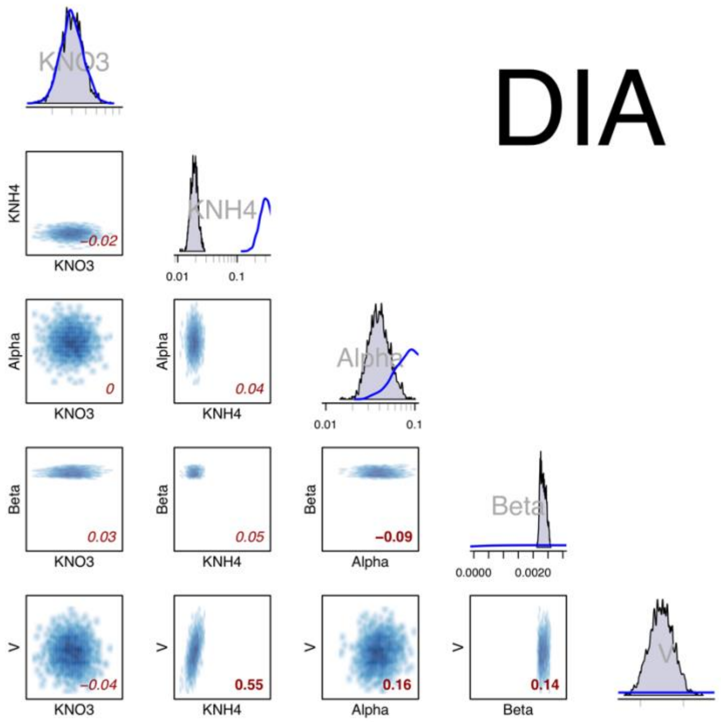
DIA
KNO3
KNH4
Alpha
Beta
V
-0.02
0
0.04
0.03
0.05
-0.09
-0.04
0.55
0.16
0.14
0.01
0.1
0.0000
0.0020

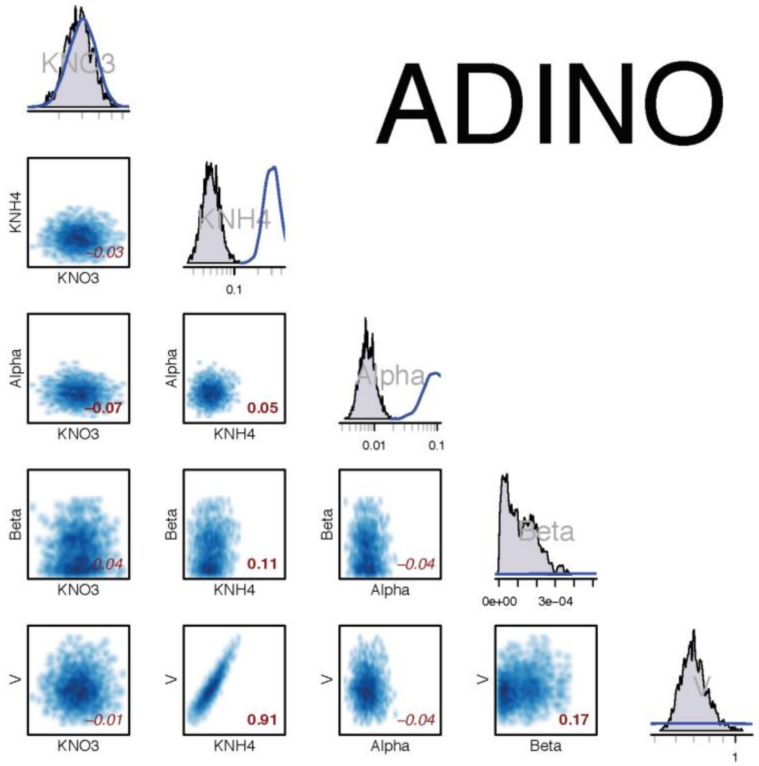
ADINO
KNO3
KNH4
Alpha
Beta
V
-0.03
-0.07
0.05
0.04
0.11
-0.04
-0.01
0.91
0.17
0.1
0.01
0e+00
3e-04
1

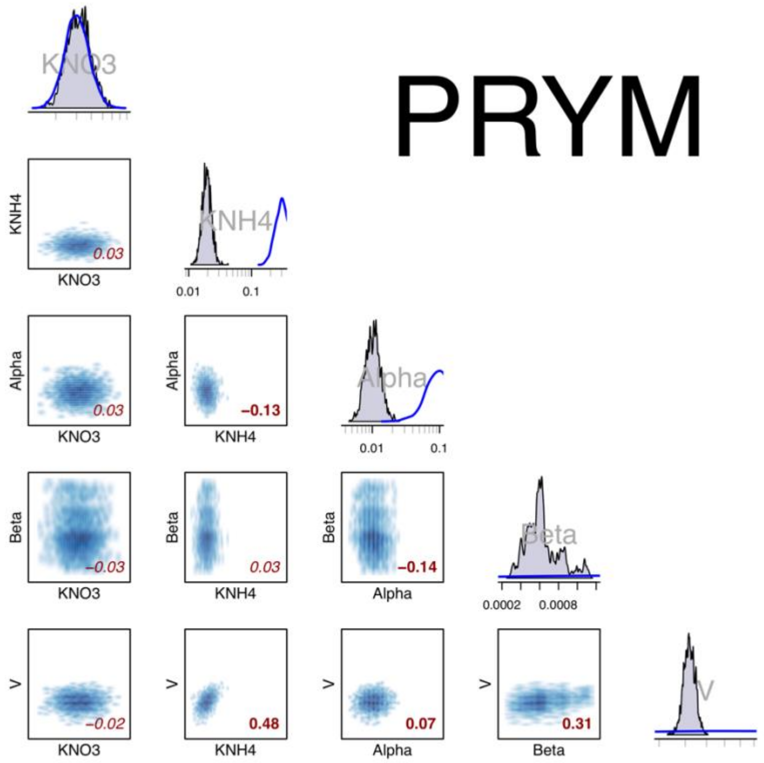


**Supplemental Figure 1.** Model parameter figures for SYN, OTHER, DIAT, ADINO and PRYM. Red value located in bottom right corner in each subplot represents correlation coefficient.

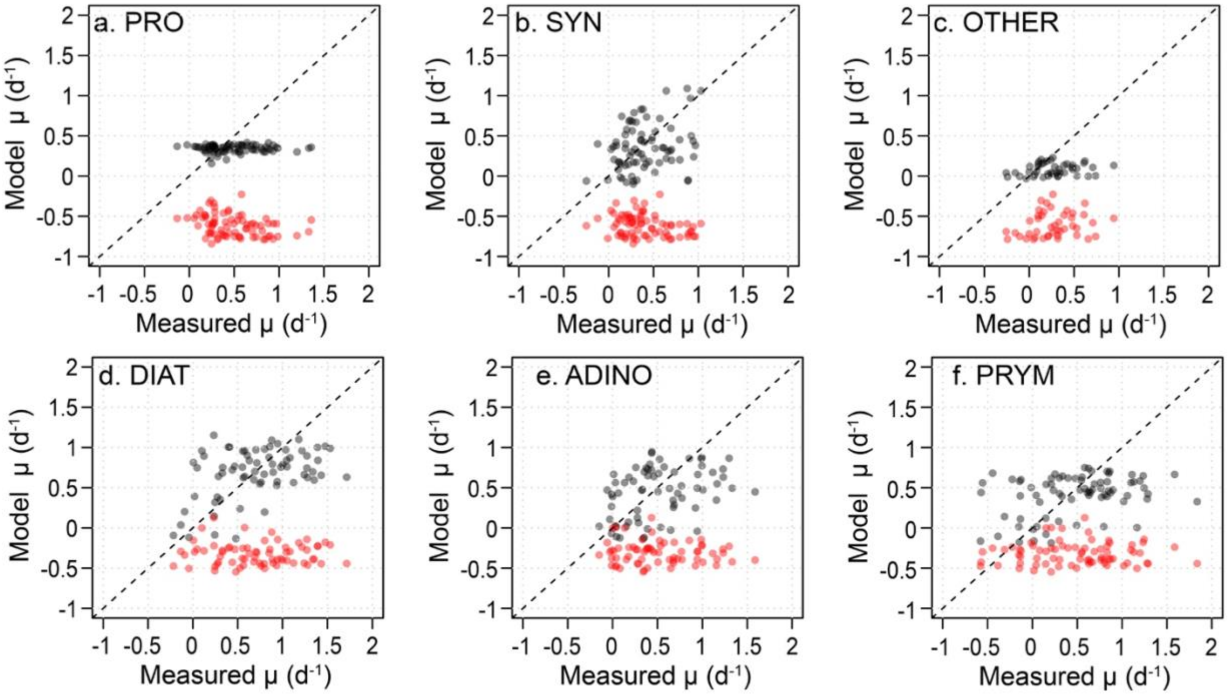


**Supplemental Figure 2.** Modeled verses measured growth rates ($d^{-1}$) for each phytoplankton group as indicated. Dashed line is 1:1 ratio. Red dots indicate NEMURO-GoM parameterization while black dots show the optimized parameters.

**Supplemental Table I.** Likelihood (Eq. 8) and prior probabilities (Eq. 9) for each observation with NEMURO-GoM parameterization (Shropshire *et al*., 2019) and the optimized model (this paper).

| **Log Likelihood or Prior Probability** | **Shropshire et al., 2019** | **Yingling et al., this paper** |
|---:|---:|---:|
| $NO_3^-$ Uptake (mmol N $m^{-3}$ $d^{-1}$) | -801 | -419 |
| NPP (mmol N $m^{-3}$ $d^{-1}$) | -31,851 | -1,615 |
| $\mu_{PRO}$ ($d^{-1}$) | -1,854 | -151 |
| $\mu_{SYN}$ ($d^{-1}$) | -1,568 | -155 |
| $\mu_{OTHER}$ ($d^{-1}$) | -844 | -187 |
| $\mu_{DIAT}$ ($d^{-1}$) | -1,614 | -232 |
| $\mu_{ADINO}$ ($d^{-1}$) | -1,066 | -221 |
| $\mu_{PRYM}$ ($d^{-1}$) | -1,252 | -351 |
| **Σ Likelihood** | **-40,748** | **-3,333** |
| **Log Prior Probability** | **-51** | **-429** |
| **Log Likelihood + Prior** | **-40,799** | **-3,762** |
